# Gene regulatory networks controlling temporal patterning, neurogenesis, and cell fate specification in the mammalian retina

**DOI:** 10.1101/2021.07.31.454200

**Authors:** Pin Lyu, Thanh Hoang, Clayton P. Santiago, Eric D. Thomas, Andrew E. Timms, Haley Appel, Megan Gimmen, Nguyet Le, Lizhi Jiang, Dong Won Kim, Siqi Chen, David Espinoza, Ariel E. Telger, Kurt Weir, Brian S. Clark, Timothy J. Cherry, Jiang Qian, Seth Blackshaw

## Abstract

Gene regulatory networks (GRNs), consisting of transcription factors and their target cis-regulatory sequences, control neurogenesis and cell fate specification in the developing central nervous system, but their organization is poorly characterized. In this study, we performed integrated single-cell RNA- and scATAC-seq analysis in both mouse and human retina to profile dynamic changes in gene expression, chromatin accessibility and transcription factor footprinting during retinal neurogenesis. We identified multiple interconnected, evolutionarily-conserved GRNs consisting of cell type-specific transcription factors that both activate expression of genes within their own network and often inhibit expression of genes in other networks. These GRNs control state transitions within primary retinal progenitors that underlie temporal patterning, regulate the transition from primary to neurogenic progenitors, and drive specification of each major retinal cell type. We confirmed the prediction of this analysis that the NFI transcription factors *Nfia*, *Nfib*, and *Nfix* selectively activate expression of genes that promote late-stage temporal identity in primary retinal progenitors. We also used GRNs to identify additional transcription factors that promote (*Insm1/2*) and inhibit (*Tbx3*, *Tcf7l1*/*2*) rod photoreceptor specification in postnatal retina. This study provides an inventory of cis- and trans-acting factors that control retinal development, identifies transcription factors that control the temporal identity of retinal progenitors and cell fate specification, and will potentially guide cell-based therapies aimed at replacing retinal neurons lost due to disease.

## Introduction

The central nervous system is highly complex, and consists of many functionally and molecularly distinct cell types, which are generated in discrete though often overlapping temporal windows (Holguera and Desplan, 2018; Oberst et al., 2019; Paridaen and Huttner, 2014). In both vertebrates and invertebrates, temporal patterning is controlled intrinsically, by dynamically regulated expression of transcription factors, which in turn regulate the ability of neural progenitors to proliferate and generate specific cell types (Cayouette et al., 2003; Doe, 2017; Rossi et al., 2021; Thor, 2017). Multiple individual transcription factors that control temporal patterning in both Drosophila (Bayraktar and Doe, 2013; Erclik et al., 2017; Konstantinides et al., 2021) and mammalian (Sagner et al., 2020; Telley et al., 2019) neural progenitors, and large-scale gene expression analysis of the developing brain has identified many other dynamically expressed transcription factors (Carter et al., 2018; Manno et al.; Tiklová et al., 2019). However, the highly diversity and poor characterization of cell types in much of developing central nervous system (Zeng and Sanes, 2017) has greatly hindered progress into our understanding of the genomic targets of these transcription factors, the networks into which they are organized, and the mechanisms by which they control temporal transitions and regulate neurogenesis.

In contrast to most brain regions, the retina represents a relatively tractable system for identifying broadly applicable molecular mechanisms controlling temporal patterning and neurogenesis in the developing central nervous system. The retina is composed of seven major cell types whose generation and molecular properties are well-characterized. Specifically, retinal ganglion cells, cone photoreceptors, horizontal cells and GABAergic amacrine cells are specified during early stages of neurogenesis prior to embryonic day (E)18, while non-GABAergic amacrine cells, bipolar cells, Müller glia and most rod photoreceptors are specified at later ages (Bassett and Wallace, 2012; Cepko, 2014; Young, 1985a). Much effort has been directed towards identifying factors that control retinal cell identity (Malin and Desplan, 2021; Sanes and Zipursky, 2010). Some transcription factors have been identified that act as master regulators of retinal cell fate specification, such as *Otx2*, which promotes photoreceptor and bipolar cell fate while repressing amacrine cell specification (Ghinia Tegla et al., 2020; Nishida et al., 2003). Several recent large-scale single-cell RNA-seq (scRNA-seq) studies have comprehensively profiled gene expression in mouse, human, and zebrafish retinas across the full course of neurogenesis (Clark et al., 2019; Cowan et al., 2020; Lu et al., 2020; Xu et al., 2020). These and other studies have identified multiple transcription factors (TFs) that are selectively expressed in either early or late-stage retinal progenitor cells (RPCs). Genetic analysis has shown that several of these are required for generation of individual retinal cell types (Clark et al., 2019; Elliott et al., 2008; Javed et al., 2020; Liu et al., 2020; Mattar et al., 2015).

Despite these advances, our understanding of the detailed mechanisms by which retinal cell fate specification is regulated remains largely unclear. Gene expression data alone does not directly identify regulatory relationships between transcription factors and their target genes. It is likewise not known how transcription factors that are selectively expressed in early or late-stage RPCs regulate temporal identity. While RPCs appear to commit to specific cell fates around the time of final exit from mitosis (Cepko, 2014), the molecular mechanisms that control this process are also still unknown. Furthermore, multiple transcription factors typically interact to coordinate progenitor cell fate transitions and cell fate specification, with feedback loops in which pairs of cell or stage-specific transcription factors regulate each other either positively or negatively playing an critical role in controlling developmental transitions (Davidson, 2001; Hobert, 2008).

In particular, the organization of the gene regulatory networks (GRNs) that control retinal neurogenesis and cell fate specification currently remains unexplored at the single cell level. While several studies have globally analyzed chromatin accessibility and/or histone modifications in either whole retina (Aldiri et al., 2017; Xie et al., 2020) or purified individual developing retinal cell types (Murphy et al., 2019; Stein-O’Brien et al., 2019; Zibetti et al., 2019), the extensive heterogeneity of cell types in the developing retina limits the usefulness of these datasets. Despite the wealth of scRNA-seq data now available, we still lack a real understanding of how transcription factors interact and dynamically regulate expression of genes controlling retinal temporal patterning and cell fate specification.

In this study, we address this by generating chromatin accessibility profiles of the developing mouse retina over the full course of neurogenesis using single-cell ATAC-seq (scATAC-seq). We identify *cis*- regulatory elements and putative transcription factor binding sites from the scATAC-seq datasets, and then integrate these results with existing, age-matched scRNA-seq datasets from mouse (Clark et al., 2019), as well as newly generated scRNA-seq and scATAC-seq datasets from developing human retina (Thomas et al., 2021) to identify evolutionarily-conserved GRNs controlling key developmental transitions and cell fate specification. Within these GRNs, cell type-specific transcription factors activate and maintain expression of other transcription factors within the network while often also inhibiting, or more rarely activating, expression of transcription factors in other networks. We identify GRNs specific to neuroepithelial-like cells, early and late-stage primary and neurogenic RPCs, and all major neuronal and glial cell types of the retina. By modelling dynamic regulatory relationships between transcription factors in cell-specific GRNs, we have been able to make and validate specific predictions about their function in retinal development. For instance, we demonstrate that the NFI factors *Nfia/b/x*, which were previously shown to promote specification of late-born retinal cell types (Clark et al., 2019), directly activate expression of transcription factors selectively expressed in late-stage RPCs and Müller glia. We also identify previously uncharacterized activators (*Insm1/2*) and inhibitors (*Tbx3* and *Tcf7l2*) of rod photoreceptor specification and differentiation. This resource provides a roadmap for the research community to identify the gene regulatory networks that control retinal development.

## Results

### Single-cell ATAC-seq profiling of developing mouse retina

To comprehensively profile dynamic changes in chromatin accessibility across the full course of retinogenesis, we conducted scATAC-seq analysis using the 10x Genomics Chromium platform on dissociated cell nuclei from whole mouse retina at 11 timepoints: embryonic (E) day 11, 12, 14, 16, and 18, as well as postnatal (P) day 0, 2, 5, 8, 11 and 14 (Figure 1A), profiling a total of 108,975 cells (Figure S1A). The distribution of the size and position of accessible DNA sequences relative to annotated transcriptional start sites (TSS) were highly consistent among each of these samples (Figure S1B), demonstrating overall high quality data. We observed high overall correlations between age-matched scATAC-seq and bulk ATAC-seq retinal progenitor (RPC) samples at E11 (r=0.82) and P2 (r=0.94), while lower correlations were seen between age-mismatched E11 and P2 samples (Figure S1C) (Zibetti et al., 2019). ScATAC-seq analysis robustly detected peaks seen in bulk ATAC-seq data, and reflected temporal differences in gene expression, as shown for the bHLH factor *Hes5*, which is strongly enriched in late-stage RPCs (Furukawa et al., 2000; Hojo et al., 2000) (Figure S1D).

**Figure 1:**
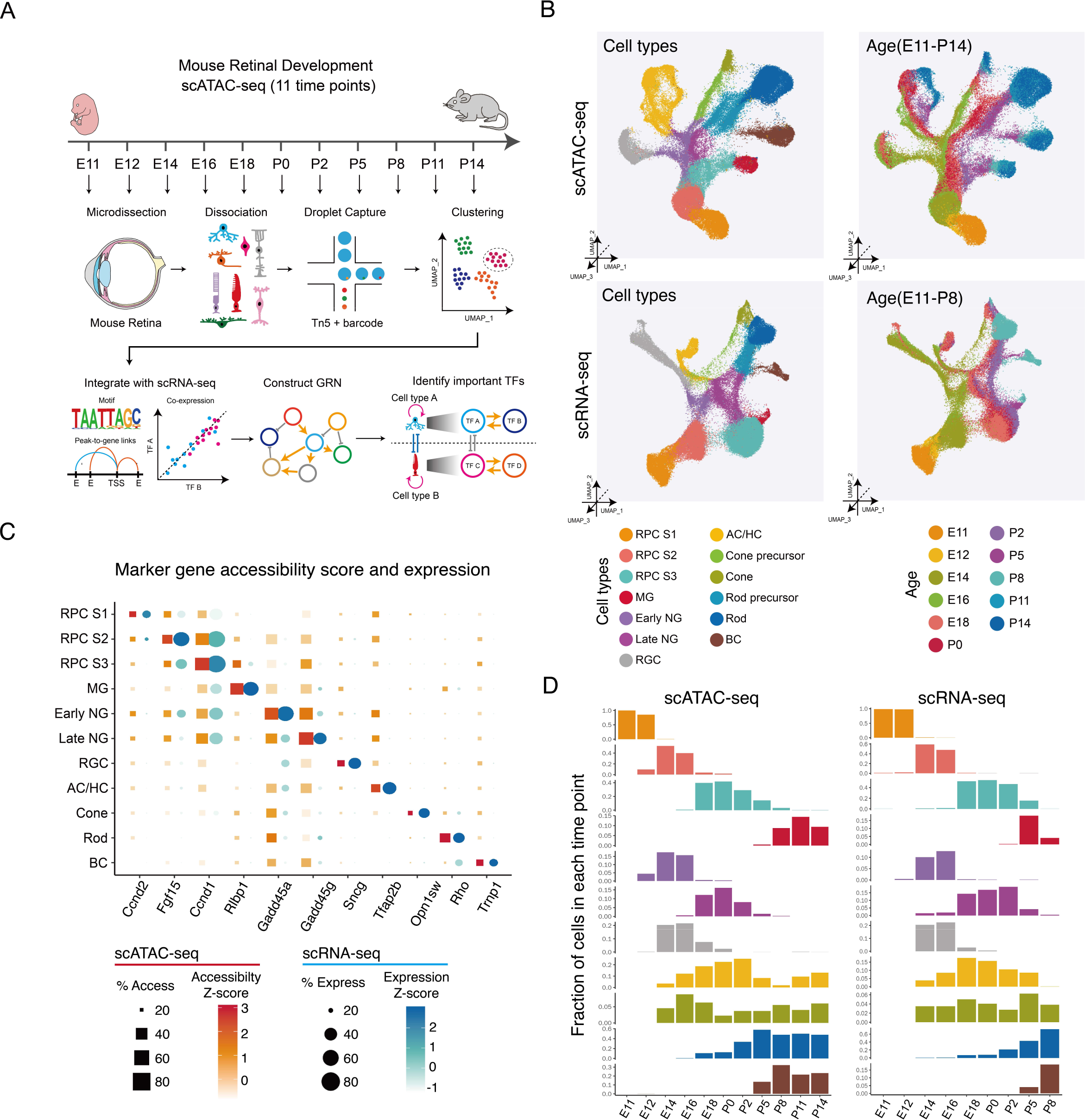
Overview of the study. (A) Schematic summary of the study. ScATAC-seq of the mouse whole retinas was performed at 11 different time points. Cell types and cell-type specific accessible chromatin regions were identified through dimensional reduction and clustering analysis. By integrating age-matched single-cell RNA-seq data with our datasets, we reconstructed gene regulatory networks (GRN) using the IReNA v2 analytic pipeline, and identified candidate regulators controlling temporal patterning and cell fate specification during the retinal development. (B) Combined UMAP projection of all mouse retinal cells profiled using scATAC-seq (top) and scRNA-seq datasets (bottom). Each point (cell) is colored by cell type (left) and age (right). (C) Examples of the expression and chromatin accessibility for selected cell type-specific genes. (D) The relative abundance of retinal cell types is similar between age-matched scATAC-seq (left panel) and scRNA-seq (right panel). Bar plots showing the fraction of cells (y-axis) at each time point of each cell type (x-axis). RPCs, retinal progenitor cells, MG, Müller glia; AC/HC, amacrine/horizontal cells; BC, bipolar cells; RGC, retinal ganglion cells; NG, neurogenic progenitor cells.

UMAP analysis was then performed on datasets obtained from each individual time point to identify individual cell types, which were annotated based on differential accessibility of a panel of well-characterized cell type-specific genes (Table ST1). UMAP analysis of scATAC-seq data showed broad similarity to age-matched scRNA-seq data (Figure 1B). Several features were observed from this analysis. First, as previously reported using scRNA-seq analysis (Clark et al., 2019), a clear distinction was seen between early-stage neuroepithelial cells (which here we term RPCs stage 1), and both early-stage and late-stage primary RPCs (RPCs stage 2 and 3 respectively), with Müller glia (MG) arising directly from late-stage primary RPCs. Distinct populations of early and late-stage neurogenic RPCs are also observed, which RNA velocity analysis indicates arise from early and late-stage primary RPCs, respectively (Melsted et al., 2021) (Figure 1B,C). Second, four major trajectories of differentiating neurons are observed: retinal ganglion cells (RGCs); amacrine and horizontal cells (AC/HC); rod and cone photoreceptors; and bipolar cells (BC), respectively (Figure 1B, Fig. S1). Third, the timing of the appearance of each retinal cell type and their relative abundance is also broadly similar between the two data sets (Figure 1D, Table ST1).

Neuroepithelial cells (RPCs stage 1) dominate the E11 and E12 samples, while RGCs, cones and amacrine/horizontal cells are detected by E14. A rapid transition between early and late-stage primary and neurogenic RPCs is seen between E16 and E18, which coincides with a dramatic reduction in the relative abundance of RGCs, as previously observed using scRNA-seq analysis (Clark et al., 2019). Likewise, late-born bipolar cells and Müller glia are first observed in both datasets at P5 (Figure 1D). We observe a high overall correlation between scATAC-seq and scRNA-seq profiles of individual cell types (Figure S1E).

A previous study has conducted ATAC-seq and ChIP-seq analysis of chromatin modifications in whole mouse retina over the course of development (Aldiri et al., 2017). We investigated the extent to which open chromatin regions (OCRs) that were detected using our scATAC-seq matched genomic annotations that were defined by Hidden Markov Modeling (HMM) from age-matched bulk retinal samples (Aldiri et al., 2017). While HMM analysis showed that the great majority of genomic regions were predicted to lack regulatory potential at all ages (Aldiri et al., 2017), we found that the majority of OCRs identified in P0 and P14 retina using aggregated scATAC-seq overlapped with regions identified as active promoters or enhancers, as defined using HMM analysis of bulk ChIP-Seq and ATAC-Seq data (Figure S1F). A comparison of OCRs present in specific cell types to age-matched HMM data revealed that the most abundant cell types showed the strongest predicted regulatory potential, with OCRs found in primary and neurogenic RPCs showing particularly high regulatory potential at P0, as did rod photoreceptor OCRs at P14 (Figure S1F). However, even in rarer cell types -- such as retinal ganglion cells (RGC) at P0 or Muller glia (MG) at P14 -- at least one-third of all OCRs were predicted to show regulatory potential. These broadly reflect overall changes in retinal cell composition, and highlight the importance of the finer resolution analysis provided by scATAC-seq data. Genes that are highly specific to different mature retinal cell types -such as *Aqp4* in MG, *Tfap2b* in amacrine cells (AC), *Opn1sw* in cones, *Rho* in rods and *Cabp5* in bipolar cells (BC) -- showed expected cell type-specific patterns of chromatin accessibility at P14 (Figure S1G).

### Analysis of dynamic chromatin accessibility and transcription factor activity during mouse retinal development

We next systematically analyzed scATAC-seq data to identify cell type-specific cis-regulatory elements (Figure 2A, Table ST2). Patterns of chromatin accessibility of well-characterized cell type-specific genes generally correlate with their mRNA expression (Figure S2A), although some transcription factors specifically expressed in neurogenic RPCs such as *Atoh7* and *Olig2* also showed high levels of accessibility in early-born neurons, including RGCs, cones, and AC/HC. All cell types showed many specific peaks of chromatin accessibility, with MG (12,757 peaks) and RGC (8,576 peaks) showing the largest number, and early neurogenic RPCs showing the smallest number of peaks (1,497). Next, we defined the gene expression that is associated with these cell type-specific accessibility regions. For each cell type-specific accessible peak, we calculated the peak-gene correlation and selected their positive correlated gene expression. In total, we identified 11,203 corresponding genes for all cell types (Figure 2B, Table ST2).

**Figure 2:**
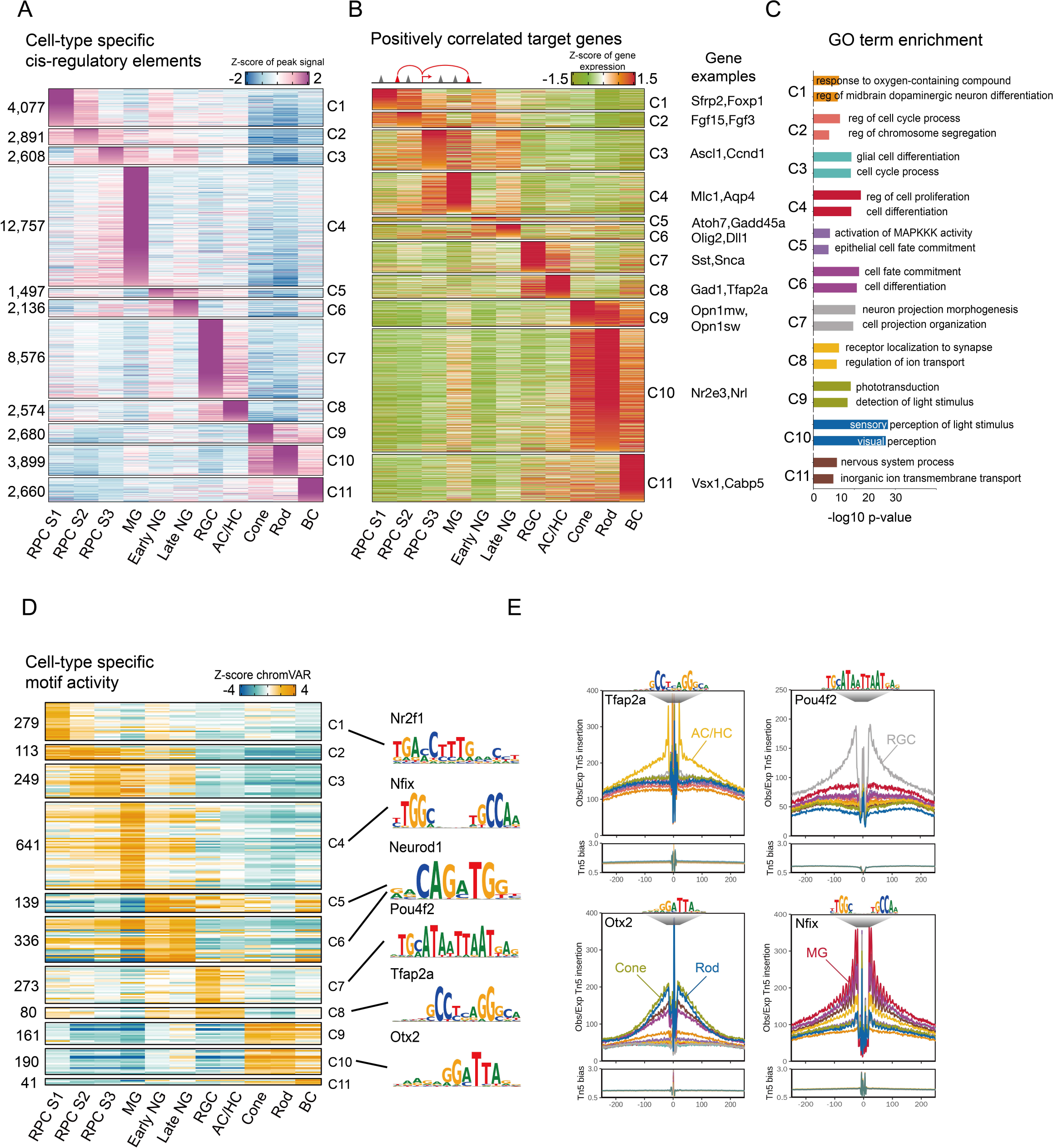
Single-cell regulatory landscape of mouse retinal development. (A) Heatmap of cell-type-specific peaks. The numbers of cell type-specific peaks are indicated on the left. Cell types are shown at the bottom. (B) Heatmaps of the expression level of positively correlated genes. Cell types are shown at the bottom of the plot. (C) Representative genes along with GO enrichment for each cluster. The X-axis indicates the - log10(P-value) of the GO term. (D) Heatmap of the chromVAR z-score for cell-type-specific motifs. The number of motifs in each cell type are indicated on the left. Cell types are indicated at the bottom. Representative motif logos are shown on the right. (E) Examples of TF footprint profiles for *Tfap2a*, *Pou4f2*, *Otx2* and *Nfix* in the indicated scATAC-seq clusters. Tn5 insertion tracks are shown below. RPCs, retinal progenitor cells, MG, Müller glia; AC/HC, amacrine/horizontal cells; BC, bipolar cells; RGC, retinal ganglion cells; NG, neurogenic progenitor cells

These include *Nrl* and *Nr2e3* in rods, *Mlc1* and *Aqp4* in Müller glia, and *Sfrp2* and *Foxp1* in stage 1+2 RPCs. Gene Ontology (GO) analysis reveals that genes associated with progenitor-specific differentially accessible regions (DARs) are often involved in cell cycle regulation; genes associated with rod and cone-specific DARs are involved in phototransduction and visual cycle; and genes associated with neurogenic RPC-specific DARs are involved in regulation of development (Figure 2C, Table ST2).

We then analyzed the activity of transcription factors that could potentially interact with the DARs. We measured the gain or loss of global chromatin accessibility in DARs containing individual TF motifs by using chromVAR software (Figure 2D, Table ST2) (Schep et al., 2017). Many transcription factors showed some degree of cell type-specificity, with the number of cell type-specific motifs ranging from 641 in Müller glia to 41 in bipolar cells (Table ST2). We further validated chromVAR scores by using footprinting analysis for known transcription factor markers that are selectively expressed in specific retinal cell types. These include *Tfap2a* motifs associated with AC/HC-specific footprints; *Pou4f2* motifs associated with RGC-specific footprints; *Otx2* motifs that are associated with photoreceptor-specific footprints; and *Nfix* motifs associated with Müller glia-specific footprints (Figure 2E, S2B). Integrated scRNA-seq and scATAC-seq analysis can thus reliably identify targets of known cell type-specific TFs in mice.

### Comparison of mouse and human scATAC-seq data reveals evolutionary-conserved regulatory elements and motif activities

To identify evolutionarily conserved regulatory elements and transcription factors controlling retinal neurogenesis and cell fate specification, we compared our mouse datasets to scATAC-seq and scRNA-seq data obtained from six developmental time points from human retina, ranging from 7.5 to 19 gestational weeks (Thomas et al., 2021). As in the mouse, UMAP analysis identifies each of the major cell types of the developing retina (Figure S3A,B, Table ST3), and closely resembles an aggregate UMAP plot previously reported for a more extensive scRNA-seq analysis of human retinal development (Lu et al., 2020)(Figure S3C). We next identified evolutionarily-conserved cell type-specific regulatory elements for all major retinal cell types. We find that 3-15% of these elements are conserved between mouse and human, with RGCs the highest, and cones and Müller glia the lowest (Figure 3A). This may in part reflect oversampling of early timepoints in the human data, as these samples are highly enriched for RGCs and have relatively few Müller glia (Table ST3, ST4). The low level of conservation of cone-specific elements, however, likely reflects the fact that cones are the most transcriptionally divergent between mice and humans (Lu et al., 2020). Overall, 8.3% of mouse peaks and 6.4% of human peaks are evolutionarily-conserved (Figure 3B, Table ST3, ST5). No clear enrichment is observed in evolutionarily-conserved peaks relative to all peaks in either species, with the exception of evolutionarily-conserved human peaks showing greater enrichment for transcriptional start sites (TSS) (4.9% vs. 2.1%, p-value<2.2e-16).

**Figure 3:**
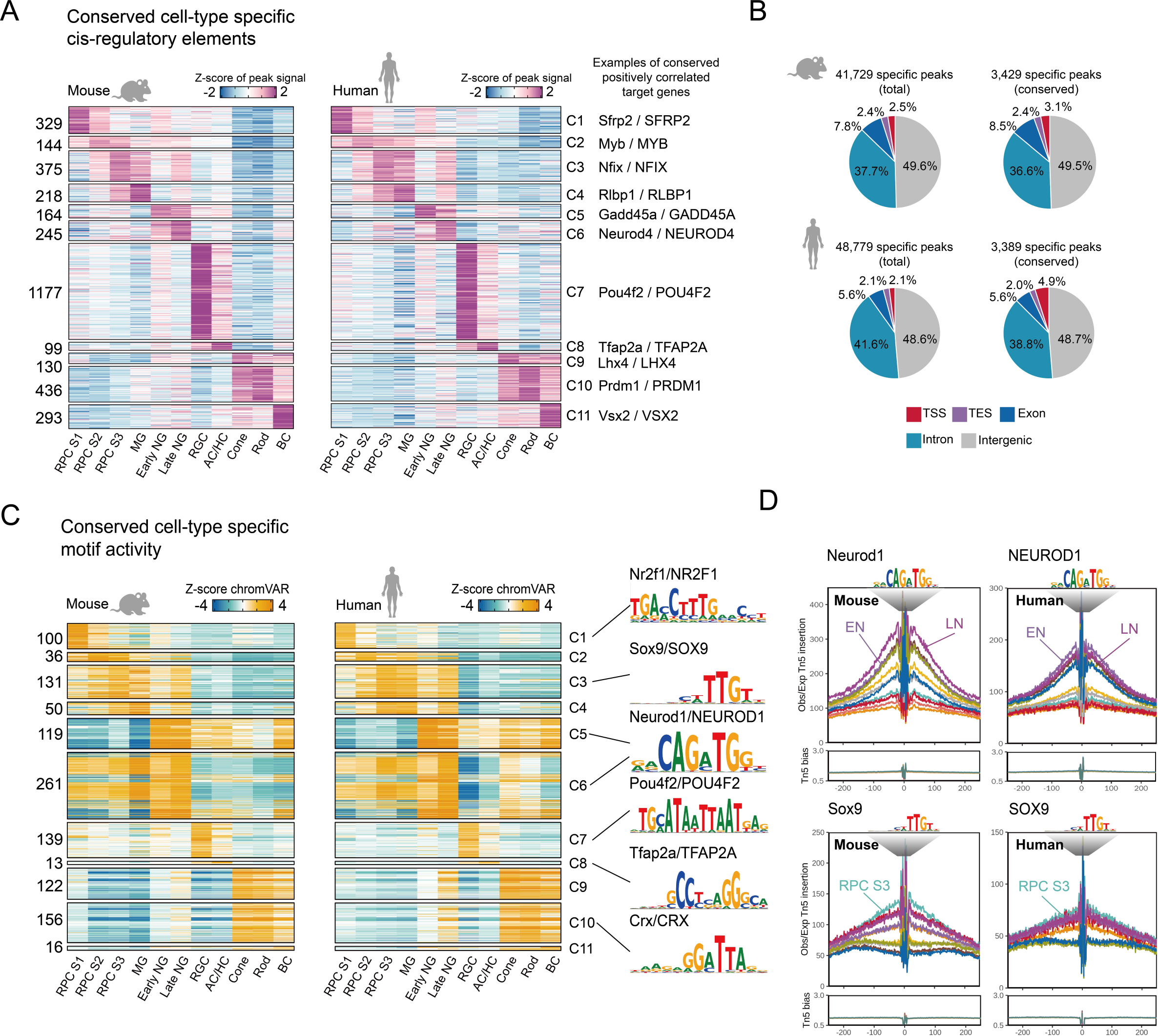
Conserved single-cell regulatory landscape in mouse and human retinal development. (A) Heatmap of evolutionarily conserved cell type-specific peaks. The numbers of peaks are indicated on the left. Cell types are shown at the bottom. Representative conserved and positively correlated genes are shown on the right. (B) Pie chart depicts the percentage of total and conserved peaks from mouse (top) and human (bottom). TSS, transcriptional start site; TES, transactional end site. (C) Heatmap of the chromVAR z-score of the conserved cell-type-specific motifs from mouse (left) and human (right). The number of motifs are indicated on the left. Cluster identities are indicated at the bottom. Representative motif logos are shown on the right. (D) TF footprint profile of *Neurod1* and *Sox9* from selected mouse and human retinal cell types. RPCs, retinal progenitor cells, MG, Müller glia; AC/HC, amacrine/horizontal cells; BC, bipolar cells; RGC, retinal ganglion cells; NG, neurogenic progenitor cells

To analyze conservation of trans-acting factors regulating cell type-specific gene expression, we compared cell type-specific active motifs in mouse and human cell types using chromVAR (Schep et al., 2017). As expected, we observed a much higher percentage of conserved active motifs of cell type-specific regulatory elements, with numbers of conserved cell type-specific motifs ranging from 122/161 in cones to 50/641 in Müller glia. Cell type-specific active motifs include well-characterized developmental regulators such as *Sox9*, *Neurod1*, *Pou4f2*, *Tfap2a* and *Crx* (Figure 3C, Table ST3, ST5). Similar developmental patterns of transcription factor footprinting are observed for many of these TFs, as illustrated by *Neurod1* in late neurogenic RPCs and *Sox9* in stage 3 RPCs and Müller glia (Figure 3D, Table ST3). Using the same analytic approach that was applied to mouse retina, we can also reliably identify targets of known cell type-specific TFs in developing human retina.

### Gene regulatory networks controlling temporal patterning of retinal progenitors

Since the generation of all retinal cell types is ultimately controlled by the dynamic temporal patterning of primary RPCs over the course of neurogenesis (Cepko, 2014; Zechner et al., 2020), we next set out to identify gene regulatory networks (GRNs) that potentially control this process. To identify key TFs, we first reconstructed GRNs by integrating scRNA-seq and scATAC-seq data using a modified form of the IReNA analysis pipeline (Hoang et al., 2020) (Figure S4A,B). We then extracted predicted regulatory relationships among stage-specific TFs, identifying positive feedback loops of co-expressed TFs used to maintain stage-specific identity, and negative feedback loops used to ensure mutually exclusive expression of TFs specific to different stages. We further filtered these results based on the strength of the predicted regulatory relationships, and whether the expression pattern of individual TFs was conserved between mouse and human retina (Figure 4A; S4A).

**Figure 4:**
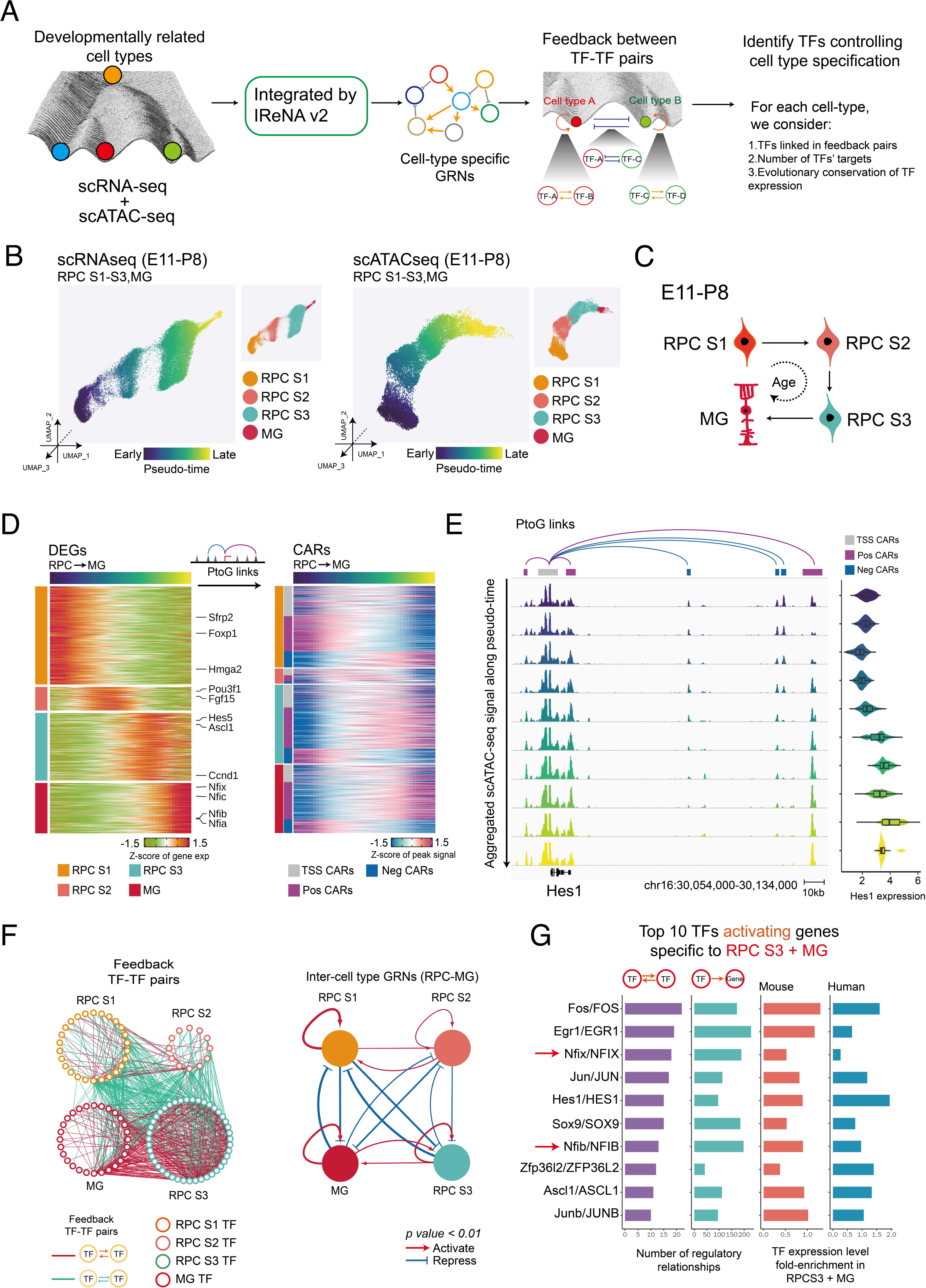
Model of gene regulatory networks controlling temporal patterning of retinal progenitors. (A) Schematic of the analytic pipeline used to identify TFs controlling retinal development. The role of feedback loops (double-positive and double-negative) in controlling transitions between cell states and during the retinal cell fate specification is shown in the Waddington epigenetic landscape model. (B) UMAPs of retinal progenitors from scRNA-seq (left) and scATAC-seq (right). Cells are colored by pseudotime and cell type. (C) A model for the transitions of primary retinal progenitors and Müller glia during E11-P8. (D) Heatmaps show the expression of cell type-specific DEGs (left) and their correlated accessible regions (CARs, right) across pseudotime. The left bar indicates the cell types (RPCs S1-S3, MG) and the classes of CARs (TSS, positively correlated and negatively correlated). (E) Genome track visualization of the *Hes1* locus. Each track represents the aggregated scATAC-seq signals across the RPC-MG trajectory. Inferred links of *Hes1*-associated CARs (correlated accessible regions) are shown on the top. Expression level of *Hes1* measured by scRNA-Seq across the RPC-MG trajectory is shown on the right. (F) Full network diagram showing TF pairs linked by reciprocal positive or negative regulatory relationships during the RPC-MG transition (left). Each node represents an individual cell type-specific TF. Each edge represents a statistically significant feedback relation between TF pairs. Simplified intermodular regulatory networks of retinal progenitors are shown on the right. Colored nodes represent specific cell types. Connections indicate statistically significant regulations among modules. (G) The top 10 TFs predicted to activate expression of genes specific to stage 3 RPCs, as inferred from IReNA v2 analysis (left). Bar plots show the expression levels of these TFs in mouse and human stage 3 RPCs progenitors (right).

We focused on four major cell states, including stage 1-3 RPCs and Müller glia, to investigate GRNs controlling temporal patterning of neurogenesis. We first performed pseudotime analysis for both scATAC-seq and scRNA-seq data from primary progenitors and Müller glia, similar to our previous analyses (Hoang et al., 2020; Lu et al., 2020) (Figure 4B). Our scRNA-seq and scATAC-seq data suggests that a large fraction of primary RPCs progressively transition between stages 1, 2 and 3, before eventually becoming Müller glia (Figure 4C). Pseudotime analysis of scRNA-seq identified differentially expressed genes (DEGs) during each of these stages, with *Foxp1* and *Sfrp2* enriched in stage 1 RPCs, *Fgf15* and *Pou3f1* in stage 2, *Ascl1* in stage 3, and *Nfia/b/x* and *Hes5* in Müller glia (Figure 4D), broadly matching previously reported results (Clark et al., 2019). ScATAC-seq data was then used to identify correlated accessible regions (CARs), which are associated with these DEGs (Figure 4D, Table ST5). CARs include the accessible peaks near transcription start sites (TSS) and distal accessible peaks (regions < 100kb from the TSS) that are either positively or negatively correlated with the DEGs (Table ST5; Figure S4B). Dynamic regulation of both positive and negative correlated distal elements can be clearly seen in the case of *Hes1*, which is most highly expressed in S3 RPCs and Müller glia (Figure 4E). At the *Hes1* locus, one distal and two proximal positively correlated distal elements show increased accessibility across pseudotime, while three negatively correlated elements show decreased accessibility. Accessibility at the TSS, however, does not change (Figure 4E).

We next infer patterns of TF binding by integrating TF expression patterns identified using scRNA-seq with footprinting in CARs identified by scATAC-seq (Figure S4B). This allowed us to identify TF-TF regulatory relationships among each of the four cell states (Figure 4F, Table ST6). Many state-specific TFs were connected in positive feedback loops that may maintain expression of state-specific TFs, while also repressing TFs specific to other cell states. Each cell state possessed a self-activating GRN, with the stage 1 and 3 RPCs and Müller glia-specific networks predicted to be strong and the stage 2 RPC network relatively weak. GRNs specific to stage 1 and 2 RPCs showed both positive and negative regulatory relationships, with positive regulatory relationships slightly dominating. A similar situation was observed with stage 3 RPCs and Müller glia, although positive regulatory relationships were relatively stronger.

Strong negative regulatory relationships were observed between stage 1/2 RPCs and stage 3 RPCs/Müller glia, respectively (Figure 4F).

### Nfia/b/x promote late-stage RPC temporal identity

We next predicted the top TFs that play an essential role in controlling RPC/Müller glia temporal identity by integrating the diverse information, including gene regulatory relationships, gene expression specificity and evolutionary conservation of gene expression patterns (Table ST6). Notably, NFI family members were among the top TFs predicted to activate expression of TFs specific to stage 3 RPCs and Müller glia (Figure 4G). Previous genetic data suggests that NFI factors *Nfia*, *Nfib* and *Nfix* play a central role in both controlling temporal identity in retinal progenitors and formation of late-born retinal cell types (Clark et al., 2019). However, it is not known whether *Nfia/b/x* are necessary to initiate or maintain activation of genes in the late-stage RPCs. Likewise, while *NFIA/B/X* overexpression in late-stage RPCs promotes cell cycle exit and generation of late-born cell types (Clark et al., 2019), it has not yet been determined whether their misexpression in early-stage RPCs is sufficient to confer a late-stage temporal identity, and to allow activation of genes specific to late-stage RPCs. To address these questions, we used *ex vivo* electroporation to overexpress human homologues of *NFIA/B/X* in E14 retina and profiled changes in gene expression and chromatin accessibility in primary RPCs at E16 and P0 using scRNA-seq and scATAC-seq, while performing similar analysis in P2 and P14 *Nfia/b/x* cKO retina (Figure 5A, S5A).

**Figure 5:**
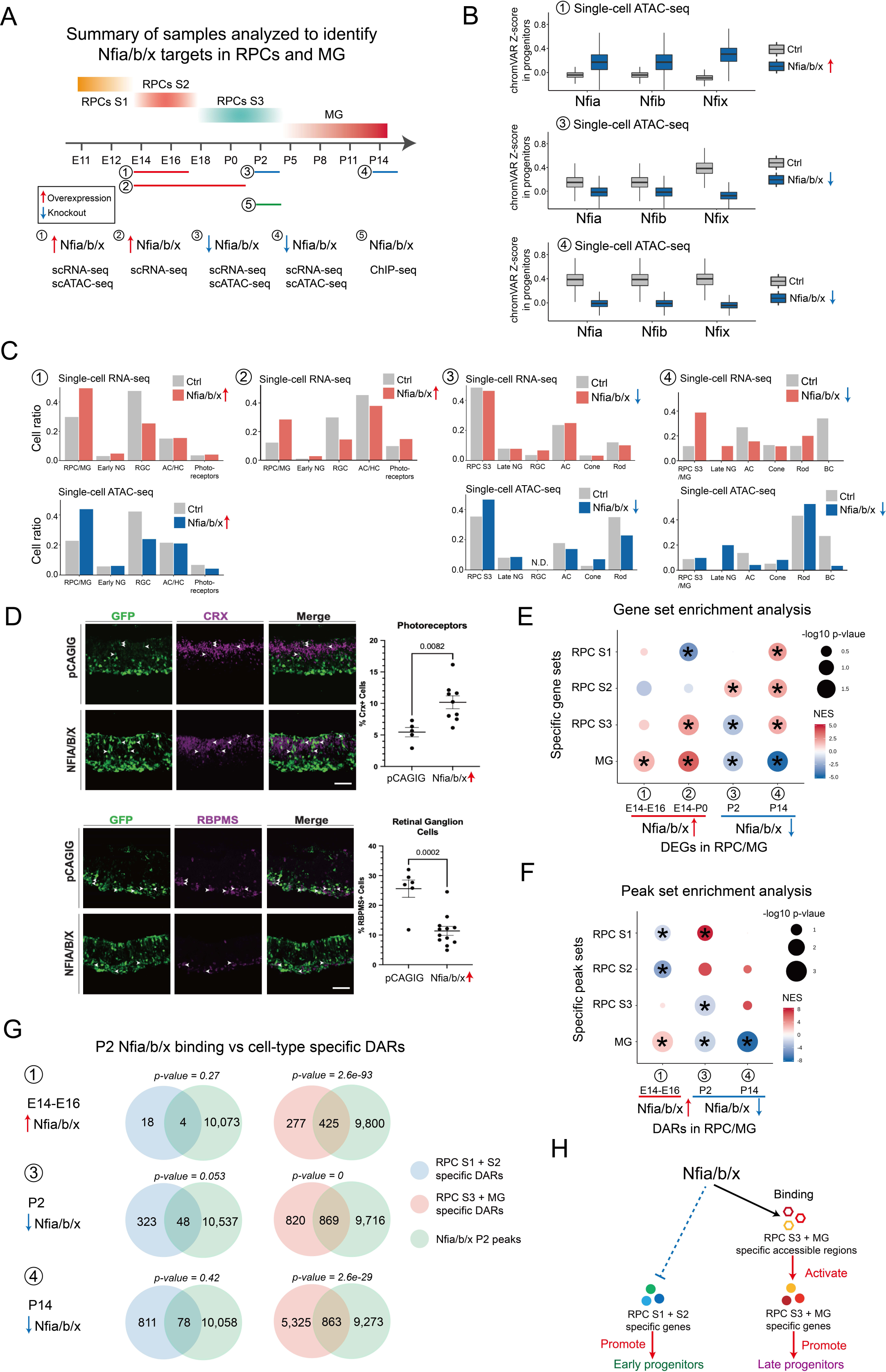
Nfia/b/x promote late-stage temporal identity in retinal progenitors. (A) Overview of experimental design to characterize the function of *Nfia/b/x* in retinal progenitor (B) Boxplots showing the changes in the *Nfia/b/x* motif enrichment in retinal progenitors following overexpression or knockout of *Nfia/b/x*. Bars are colored by genotype. (C) Bar plots showing the fraction of each retinal cell type by ages and genotypes (Top: scRNA-seq datasets. Bottom: scATAC-seq datasets) (D) Immunostaining showing fewer RGCs and more photoreceptors at P0 following *NFIA/B/X* overexpression at E14 retinal explants. The fractions of RGCs and photoreceptors are shown on the right. Scale bars = 20 μm. (E) Dot plot showing the gene set enrichment results for DEGs enriched in early and late-stage RPCs/MG following overexpression or knockout of *Nfia/b/x*. (F) Dot plot showing the peak set enrichment results for DARs enriched in early and late-stage RPCs/MG following overexpression or knockout of *Nfia/b/x*. (G) Venn diagrams showing the overlap between direct Nfia/b/x binding regions identified using ChIP-Seq and cell-type-specific DARs. The p-value on the top of each Venn diagram indicates the significance of their overlap using the hypergeometric test. (H) Summary of Nfia/b/x action during the transition from early to late-stage RPCs.

ScATAC-seq analysis demonstrates that *NFIA/B/X* overexpression induces a widespread increase in chromatin accessibility associated with sequences containing the consensus NFI motif, as has been reported in non-neuronal cells (Denny et al., 2016), while loss of function of *Nfia/b/x* leads to a loss of accessibility at these sites (Figure 5B). Relative to retinas electroporated with a control GFP plasmid, E16 retinas overexpressing *NFIA/B/X* show a reduced fraction of RGCs and an increased fraction of primary RPCs by both scRNA-seq and scATAC-seq (Figure 5C). At P0, scRNA-seq analysis and immunohistochemistry show that *NFIA/B/X*-overexpressing retinas have an increased fraction of primary RPCs and photoreceptors, and a reduced fraction of RGCs, as compared with the control (Figure 5C,D).

ScATAC-seq analysis of P2 *Nfia/b/x* cKO retinas shows an increased fraction of RPCs, along with a reduction in rod photoreceptors and AC/HC neurons (Figure 5C). In P14 *Nfia/b/x* cKO retina, substantially more RPCs are detected, while bipolar neurons are virtually absent (Figure 5C).

To determine whether gain or loss of function of *Nfia/b/x* altered expression of genes and patterns of chromatin accessibility specific to any of the three different RPC states or Müller glia, we performed gene/peak set enrichment analysis (GSEA, PSEA). We observed that overexpression of *NFIA/B/X* led to an upregulation of Müller glia-enriched genes in primary RPCs by E16 (Figure 5E, S5B, Table ST8). By P0, this effect was more pronounced, with upregulation of stage 3 RPC-enriched genes also seen. Furthermore, stage 1 RPC-enriched genes were strongly down-regulated relative to GFP controls (Figure 5E, S5B). The opposite pattern was observed in *Nfia/b/x* cKO retina, with stage 3 RPC and Müller glia-enriched genes downregulated and stage 2 RPC-enriched genes upregulated at P2. By P14, downregulation of Müller glia-enriched genes and upregulation of stage 1 and stage 2 RPC-enriched genes were more prominent (Figure 5E, S5B). Changes in patterns of chromatin accessibility closely matched those of gene expression, with NFIA/B/X overexpression inducing RPCs to adopt a state resembling Müller glia, and loss of function inducing a state that resembles stage 1 and 2 RPCs (Figure 5F, S5C). Motif analysis indicated that NFIA/B/X motifs were highly enriched in DARs that were upregulated following *NFIA/B/X* overexpression and downregulated in *Nfia/b/x* cKO retina (Figure S5C). These data demonstrate that NFI factors directly regulate temporal patterning in RPCs.

To identify genes that are directly regulated by NFIA/B/X, we performed ChIP-seq analysis on P2 wild-type retina using antibodies that recognize all three NFI factors (Table ST9). We identified 13,680 NFIA/B/X ChiP-seq peaks (Figure S5D), the majority of which (83.9%) are located in open chromatin (Figure S5E). These peaks are primarily distributed in intergenic and intronic regions. Compared to all accessible regions, NFI factor binding sites are enriched in accessible regions associated with genes specific to stage 3 RPCs (Figure S5E). We then asked whether the DARs identified following gain and loss of function of *Nfia/b/x* were direct targets for NFIA/B/X factors. By comparing NFIA/B/X ChIP-seq peaks with these DARs, we found that in E16 *NFIA/B/X*-overexpressing primary RPCs, 61% (425/702) of DARs specific to stage 3 RPCs and/or Müller glia were directly bound by NFIA/B/X, but only 4/22 of DARs specific to stage 1 and/or 2 RPCs were directly bound (Figure 5G). In P2 *Nfia/b/x* cKO retina, we observed that 51% (869/1689) of DARs specific to stage 3 RPCs and/or Müller glia overlapped with NFIA/B/X ChIP-seq peaks, in contrast to only 13% (48/371) DARs specific to stage 1 and/or 2 RPCs. Similar pattern was also observed in P14 *Nfia/b/x*cKO retina (Figure 5G). This result suggests that DARs specific to stage 3 RPCs and/or Müller glia could be directly and selectively regulated by the binding of NFIA/B/X at these regions. The direct regulatory effect of NFI factors was confirmed by the comparison of DEGs and NFIA/B/X ChIP-seq peaks (Figure S5F). In summary, our analysis suggests that NFI factors alter the chromatin accessible regions specific to late progenitor-specific genes by direct binding to these regions, and in turn activates expression of stage 3 RPCs and/or Müller glia genes and repress the expression of early progenitor-specific genes.

This underlies the mechanism by which NFI factors control temporal identity in retinal progenitors and formation of late-born retinal cell types.

### Gene regulatory networks controlling specification of retinal neurons

We applied the same approach to identify GRNs controlling neurogenesis and the specification of individual types of neurons. To do this, we generated three combined datasets, corresponding to early, intermediate and late stages of retinal neurogenesis, identifying both DEGs and CARs that are selectively active as cells adopt different identities (Table ST10). The early dataset consists of E14 and E16 time points, which included all early-stage neurogenic progenitors, as well as differentiating retinal ganglion cells, cone photoreceptors, and early-born amacrine and horizontal cells (Figure 6A). The intermediate dataset consists of E18, P0 and P2 time points, which included late-stage neurogenic progenitors, as well as differentiating rod photoreceptors and late-born amacrine cells (Figure S6A). The late dataset consists of P5 and P8 time points, and included late-stage neurogenic progenitors, and differentiating rod and bipolar cells (Figure S6E).

**Figure 6:**
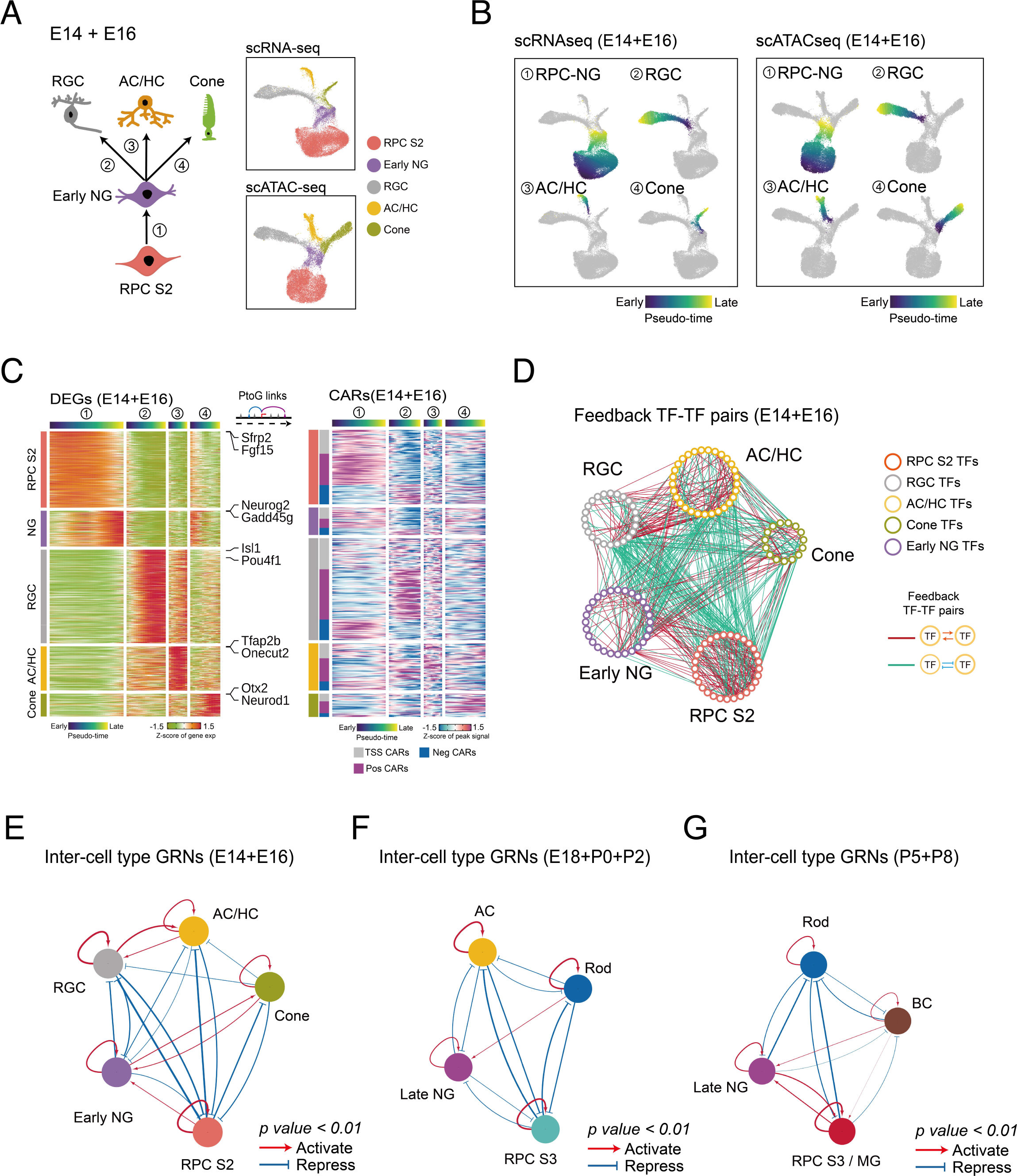
Model of gene regulatory networks controlling specification of retinal neuronal cell types. (A) A diagram showing the development of early-born retina cell types (left). UMAPs of scRNA-seq and scATAC-seq data from E14-E16 retina (right). Color indicates cell type. (B) UMAPs showing the trajectories constructed from scRNA-seq and scATAC-seq at E14-E16. Color indicates pseudotime state. (C) Heatmaps showing the expression of cell-type-specific DEGs (left) and the accessibility of their corresponding CARs (right) along differentiation trajectories. The top bars are colored by pseudotime state for each trajectory. The left bar indicates cell type and the classes of CARs. (D) Networks showing feedback relationships among TF pairs at E14-E16. Each node represents an individual cell type-specific TF. Each edge represents a positive or negative feedback regulatory relationship between TF pairs. (E-G) Simplified intermodular gene regulatory networks of RPCs and neurons at different stages (E: early-stage, F: intermediate-stage, G: late-stage). Colored nodes represent cell types. Connections indicate statistically significant regulatory relationships among GRNs specific to each cell type.

Pseudotime analysis of the early dataset revealed four major developmental trajectories.

Specifically, these were the transition from stage 2 primary RPCs to early neurogenic RPCs; and the transitions from early neurogenic RPCs to retinal ganglion cells, to amacrine and horizontal cells, and to cone photoreceptors (Figure 6B). To determine the GRNs controlling these transitions, we identified both DEGs and CARs for each developmental trajectory (Figure 6C), and inferred putative regulatory relationships among cell specific TFs (Figure 6D,E), as we had previously done for primary RPCs and Müller glia. A similar analysis was performed for intermediate (Figure S6B-D) and late (Figure S6F-H) stages of neurogenesis. This identified many TFs that were predicted to selectively activate or repress genes specific to individual cell types (Table ST11,12).

We observe many similarities between the predicted GRNs controlling neuronal cell fate specification and networks controlling state changes in primary RPCs (Fig. 4F). Cell type identity is maintained by strong positive regulatory relationships among cell type-specific TFs (Figure 6E-G; Table ST11). Regulatory relationships among different cell types often contain both positive and negative components. GRNs specific to primary RPCs and all neuronal subtypes are connected by many, almost exclusively negative regulatory relationships, while GRNs specific to most neuronal cell types are generally connected by fewer negative regulatory relationships, with positive regulatory relationships predominantly connecting GRNs of some transcriptionally similar cell types, such as RGC and AC/HC. The regulatory relationship between neurogenic and primary RPCs is more dynamic and complex. It is weakly positive at early stages of neurogenesis, weakly negative during intermediate stages, and strongly positive at late stages. This shift may reflect the fact that a rapid increase in the relative fraction of neurogenic RPCs relative to primary RPCs occurs during late neurogenesis (Clark et al., 2019). The regulatory relationship between neurogenic RPCs and neuronal networks is likewise dynamic. During early stages of neurogenesis, neurogenic RPC networks strongly inhibit RGC networks, weakly inhibit horizontal/early-born amacrine networks, and weakly activate cone networks (Figure 6E). At intermediate stages, they weakly inhibit late-born amacrine networks and weakly inhibit rod networks (Figure 6F). At late stages, they strongly inhibit rod networks but activate bipolar networks (Figure 6G). Notably, these regulations roughly correspond to the order in which these neuronal subtypes are generated during retinal development, with retinal ganglion cells formed first and bipolar cells last (Cepko et al., 1996)

### Identification of transcription factors controlling neurogenesis and cell fate specification in postnatal retina

Our GRN analysis predicts that many TFs act as either positive or negative regulators of neurogenesis and/or specification of individual neuronal subtypes. Many of these predicted regulatory relationships have been previously validated using genetic analysis (Fig. S7A, Table ST12). Considering TFs with the highest number of predicted regulatory relationships that are active in E18-P2 retina, for instance, we find that Otx2, Crx, Prdm1, Rax, Rorb, Nrl, and Nr2e3 are all predicted to activate expression of rod-specific genes (Akhmedov et al., 2000; Brzezinski et al., 2010; Furukawa et al., 1997; Irie et al., 2015; Jia et al., 2009; Mears et al., 2001; Nishida et al., 2003); Pax6, Tfap2a, and Tfap2b are predicted to repress photoreceptor specification (Jin et al., 2015; Remez et al., 2017); and Zfp36l1/2, Nfia, Hes1/5, Sox2/8/9, and Lhx2 are predicted to both promote RPC maintenance and inhibit rod differentiation (Bosze et al., 2020; Clark et al., 2019; Marquardt et al., 2001; de Melo et al., 2016; Muto et al., 2009; Roy et al., 2013; Taranova et al., 2006; Wall et al., 2009; Wu et al., 2020). Knockdown of *Nfib*, predicted to be one of the top activators of rod-specific genes, has also been recently shown to reduce rod-specific gene expression in human organoid cultures (Xie et al., 2020). Given the success in predicting the function of these TFs, we conducted gain- and loss of function analysis for several previously uncharacterized candidate TFs on retinal explants via electroporation and analyzed the resulting phenotypes using scRNA-seq and immunohistochemistry to determine whether other TFs with high numbers of regulatory relationships showed predicted phenotypes (Fig. 7A). We analyzed five different TFs: Insm1, Insm2, Tcf7l1,Tcf7l2 and Tbx3. Insm1 and Insm2 are predicted to activate genes specific to neurogenic progenitors and rod photoreceptors. In contrast,Tcf7l1/2 are predicted to inhibit neurogenesis and promote a stage 3 primary RPC/Müller glia identity. Lastly, Tbx3 is predicted to inhibit rod specification while promoting amacrine formation (Figure S7A-C).

**Figure 7:**
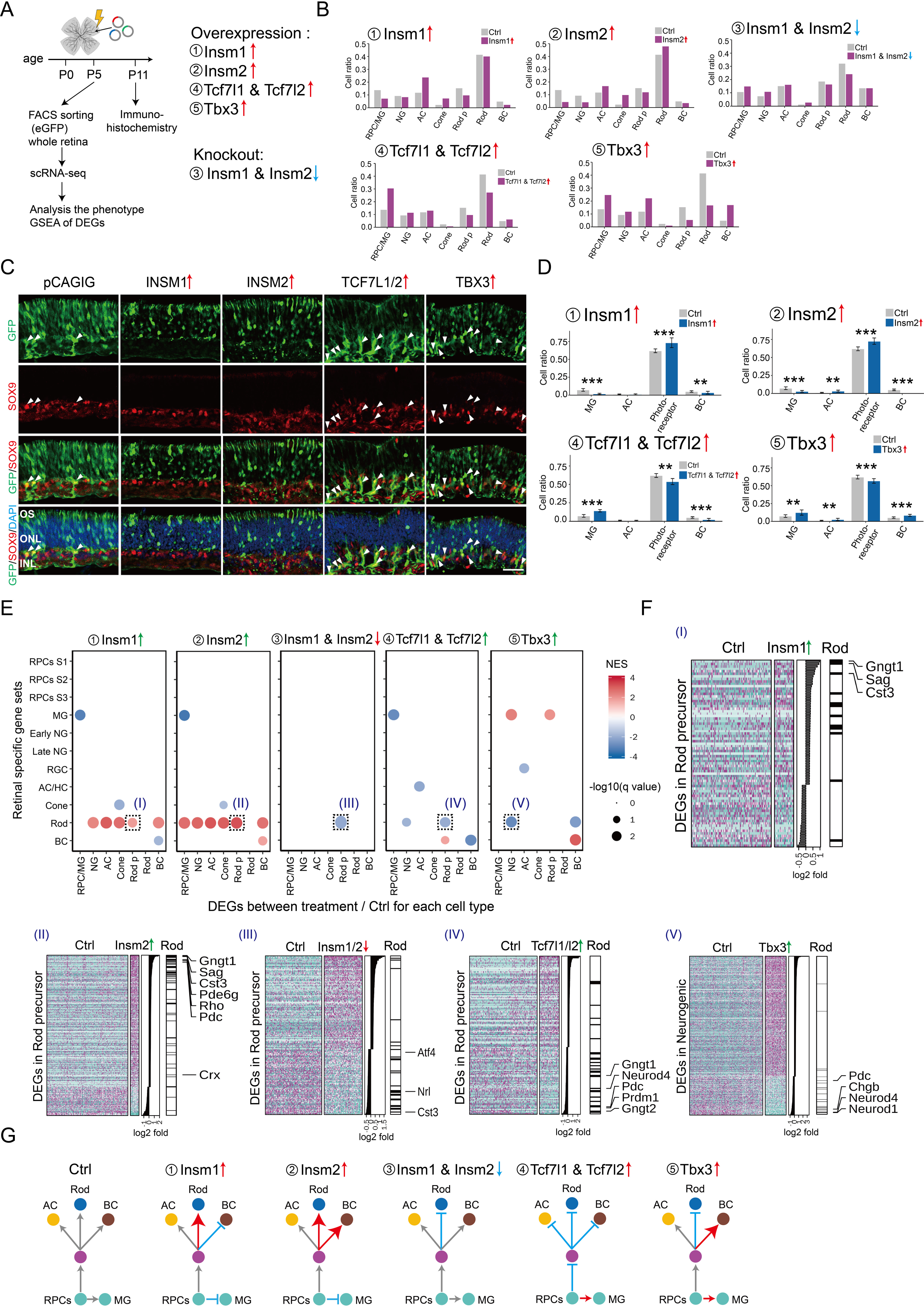
Identification of transcription factors controlling cell fate specification in postnatal retina. (A) A schematic diagram for gain-and loss-of-function analysis of candidate TFs in postnatal mouse retina explants. (B) Bar plots showing the fraction of each cell type at P5 as measured by scRNA-Seq analysis of FACS-isolated GFP-positive cells for each treatment condition. (C,D) Immunohistochemistry and quantification of Müller glia (SOX9 positive) and photoreceptors (GFP-positive in the ONL layer) in P11 retina explants in control and overexpression of INSM1, INSM2, TCF7L1/2 and TBX3. Arrow heads indicate SOX9/GFP double positive cells. Error bars indicate standard deviation. **P < 0.05, ***P < 0.001. ONL, outer nuclear layer; INL, inner nuclear layer; OS, outer segment. (E) Gene set enrichment analysis of the DEGs from each cell type in each experiment. GSEA was performed with the cell-type-specific gene sets obtained from the combined scRNA-seq datasets (E11-P8). Only significant enrichment results (p < 0.05) are shown in the dot plot. Each dot was colored by NES and sized by -log(p-value). The x-axis indicates the cell type where DEGs are calculated. The y-axis indicates the specific gene sets used in the analysis. (F) Examples of GSEA results from (E). Heatmaps show the DEGs used in the GSEA analysis, with DEGs ranked by log2 fold change, as shown in the middle panel. The right annotation shows the distribution of significantly enriched gene sets among the DEGs. Representative cell-type-specific genes are also labeled. (G) Summary of observed phenotypes. RPCs, retinal progenitor cells, MG, Müller glia; AC/HC, amacrine/horizontal cells; BC, bipolar cells; RGC, retinal ganglion cells; NG, neurogenic progenitor cells.

Overexpression of either *Insm1* or *Insm2* at P0 leads to a dramatic reduction in the relative fraction of primary RPC/Müller glia cells and an increase in the fraction of amacrine cells and cone photoreceptors at P5, as measured by scRNA-Seq analysis of FACS-isolated, GFP-positive electroporated cells (Figure 7B). *Insm2* overexpression also leads to a modest increase in the fraction of rods. Immunohistochemical analysis of P11 retinas shows that *Insm1 or Insm2* overexpression significantly increases the fraction of GFP+ cells in the photoreceptor layer, and leads to a corresponding decrease in the Müller glia fraction (Fig. 7C,D). Both *Insm1* and *Insm2* strongly activate expression of rod-specific genes in all other cell types (Figure 7E). *Insm1* and *Insm2* both accelerate the normal developmental increase in the expression of rod-specific genes in rod precursors, such as *Gngt1*, *Sag* and *Rho* (Figure 7F). In contrast, somatic CRISPR-mediated loss of function of *Insm1/2* leads to a reduction in the total number of rods, and increase in the fraction of primary and neurogenic RPCs, as well as a reduction in the expression of rod-specific genes in rod precursors, notably including *Nrl* (Figure 7B,E; Table ST13), although no statistically significant changes in cell composition were observed following loss of function of *Insm1/2* at P11 (data not shown). While *insm1a* has been reported to be required for rod differentiation in zebrafish (Forbes-Osborne et al., 2013), our data demonstrates that *Insm1/2* act to promote retinal neurogenesis, rod photoreceptor specification and rod-specific gene expression.

*Tcf7l1/2* are highly expressed in RPC/Müller glia and predicted to inhibit rod specification while maintaining RPC identity (Figure S7A,B). ScRNA-seq analysis of P5 GFP-positive cells overexpressing *TCF7L1/2* revealed a substantial increase in the fraction of stage 3 RPC/Müller glia cells, with a corresponding reduction in the fraction of rods (Fig. 7B). Immunohistochemical analysis of P11 retinas overexpressing *TCF7L1/2* leads to a reduction in relative fraction of rod photoreceptors and bipolar cells, and a corresponding increase in the fraction of Müller glia (Fig. 7C,D). In rod precursors, *Tcf7l1/2* strongly inhibits expression of both rod-specific genes such as *Gngt1* and *Pdc*, as well as transcription factors that promote rod specification such as *Prdm1* (Fig. 7E,F; Table ST14).

Finally, *Tbx3* overexpression reveals a reduction in the relative fraction of rod photoreceptors at P5, along with a corresponding increase in the fraction of not only amacrine cells, but also all other inner retinal cell types, including stage 3 RPC/Müller glia and bipolar cells (Figure 7B). Immunohistochemical analysis of retinas overexpressing *Tbx3* revealed that these changes in cell composition are maintained at P11 (Fig. 7C). *Tbx3* overexpression also leads to a reduction in expression of *Neurod1* in neurogenic progenitors, a transcription factor that promotes rod differentiation (Akagi et al., 2004) (Fig. 7E,F; Table ST14). These results, which are summarized in Figure 7G, validate our putative GRNs, and demonstrate that multiplexed scRNA-seq analysis can be scaled to analyze the function of major transcriptional regulators of retinal development.

## Discussion

This study provides a comprehensive picture of the cellular-level landscape of dynamic chromatin accessibility over the full course of retinal development, and provides both a map of candidate cis-regulatory elements and of transcription factor binding patterns. By integrating scRNA-seq data with scATAC-seq data, we reconstructed transcriptional regulatory networks that control all aspects of retinal development, including the transition between different stages of primary retinal progenitors, the transition from primary to neurogenic progenitors, and the acquisition of terminal cellular identity. We observe similarities between the retinal and other systems in both the general mechanisms and in the specific genes that control this process. For instance, much like in Drosophila, transcription factors that control these transitions act to both promote expression of genes specific to individual cell states while inhibiting expression of genes specific to earlier, later or alternative states (Doe, 2017; Rossi et al., 2021). Furthermore, several individual genes -- including *Nfia*, *Nfib*, and *Pou2f2* -- are confirmed or predicted to control temporal patterning in both retina and spinal cord, (Sagner et al., 2020).

Our work fills an important gap in our understanding of how temporal patterning is controlled. While several recent studies have used scATAC-seq to identify active transcription factor motifs in specific cell types of the developing brain (Domcke et al., 2020; Kim et al.; Sarropoulos et al., 2021), only one recent study has systematically integrated this data to identify GRNs controlling neurogenesis and specification of major neuronal and glial cell types (Di Bella et al., 2021). Moreover, while previous studies have used ATAC-seq, ChIP-seq, and HiC analysis to profile changes in both chromatin accessibility and conformation, as well as histone modifications over the course of retinal development, the information in these datasets has been limited by the analysis of highly heterogeneous cell populations (Aldiri et al., 2017; Norrie et al., 2019; Xie et al., 2020). Our scATAC-seq data allow us to visualize developmentally dynamic changes in chromatin accessibility in each major retinal cell type. Furthermore, direct comparison of stage-matched mouse and human datasets demonstrates substantial conservations of cell type-specific patterns for both gene expression and chromatin accessibility, and allows efficient identification of both evolutionary-conserved and species-specific components of GRNs that control retinal neurogenesis and cell fate specification.

Neurogenic RPCs selectively express neurogenic bHLH factors such as *Atoh7*, *Ascl1*, *Olig2* and *Neurog2* (Brown et al., 1998; Brzezinski et al., 2011; Hafler et al., 2012; Hufnagel et al., 2010; Ma and Wang, 2012). Genetic lineage analysis shows that they have a limited ability to proliferate and will undergo either asymmetric or terminal neurogenic divisions. In addition, expression of many transcription factors that are master regulators of retinal cell identity -- such as *Otx2*, *Onecut1* and *Onecut2*, and *Foxn4* -- are first detected in neurogenic RPCs (Emerson et al., 2013; Liu et al., 2020; Muranishi et al., 2011; Wang et al., 2014). Taken together, this suggests that retinal neuronal identity is specified during the neurogenic RPC stage, and that identifying gene regulatory networks controlling the transition from primary to neurogenic RPCs, and from neurogenic RPCs to postmitotic neurons, will help to understand retinal cell fate specification.

Integrated scRNA-seq and scATAC-seq analysis reveals the highly redundant and complex patterns of gene regulatory relationships that maintain each of the cellular states, and ensures that developmental processes remain consistent and robust in the face of a variety of environmental perturbations. This may explain the observation that genetic disruption of individual cis-regulatory elements typically results in only developmental phenotypes that are either modest, or only alter expression of a subset of the cell type-specific transcription factors regulated by these genes (Chan et al., 2020; Ghiasvand et al., 2011). We observe that GRNs controlling retinal development are highly parallel, redundant, and complex. Cell states are maintained by networks of TFs that activate expression of TFs within cell type-specific GRNs, but often also repress expression of TFs in GRNs specific to other cell states, potentially mediating rapid and irreversible transitions between different stable transcription states. Regulatory relationships among individual cell type-specific GRNs are temporally dynamic, often containing a mixture of positive and negative feedback loops, and accurately reflect observed developmental changes in the timing of retinal neurogenesis. For instance, while GRNs of primary RPCs show only weakly positive or negative regulation of GRNs specific to neurogenic RPCs between E14 and P2, they strongly activate neurogenic RPC-specific GRNs at P5 and P8, corresponding to the increase in the relative fraction of terminal neurogenic divisions at these ages (Cepko, 2014), as well as the increased fraction of neurogenic RPCs relative to primary RPCs at this age (Clark et al., 2019). Likewise, neurogenic RPC-specific GRNs most strongly activate expression of TFs in GRNs specific to the neuronal subtypes that are generated latest at each stage: cones at E14-E16, rods at E18-P2, and bipolar cells at P5-P8 (Young, 1985b). Identifying the precise mechanisms that control these dynamic changes in the organization of cell type-specific GRNs awaits more detailed functional analysis.

Transitions between cell states are driven by changes in both gene expression and chromatin accessibility. In some cases, TFs act to alter chromatin accessibility at regulatory sites associated with stage-specific genes prior to initiation of changes in gene expression (Ma et al., 2020). This is clearly seen with *Nfia/b/x*, which are enriched in late-stage RPCs and Müller glia. *Nfia/b/x* overexpression triggers increased accessibility at regulatory sites associated with genes expressed in late-stage RPCs and Müller glia, while loss of function of *Nfia/b/x* produces the opposite effect. This leads to activation of expression of these genes and, indirectly, to repression of genes specific to early-stage RPCs, ultimately inhibiting generation of early-born cell types such as RGCs and promoting rod photoreceptor specification. This establishes NFI factors as bona fide regulators of temporal patterning in RPCs, and identifies the mechanism by which they regulate changes in retinal progenitor competence.

Notably, *Nfib* is also predicted to be a major component of the GRN controlling rod photoreceptor differentiation (Fig. S7A), and directly targets multiple genes known to promote rod specification, including *Otx2*, *Prdm*1, *Mef2d,* and *Caszl1* (Andzelm et al., 2015; Brzezinski et al., 2010; Mattar et al., 2015; Nishida et al., 2003) While the loss of function of *Nfib* leads to a reduced number of rod photoreceptors in P2 retina (Fig. 5C), and a reduction in the expression of rod-specific genes (Xie et al., 2020), a substantially increased proportion of rod photoreceptors is seen at P14 relative to controls (Fig. 5C)(Clark et al., 2019). This implies that, while *Nfib* may activate genes that promote rod differentiation, it is ultimately dispensable for rod specification.

The network analysis conducted here identifies TFs that show high degrees of connectivity, and are hence likely to play more important roles in controlling development. We have tested the function of a subset of TFs predicted to either positively or negatively regulate retinal neurogenesis and rod photoreceptor differentiation in early postnatal retina, using overexpression or CRISPR-based loss of function analysis in combination with scRNA-Seq analysis and immunohistochemistry. This confirmed our prediction that *Insm1* and *Insm2* promote postnatal retinal neurogenesis and rod photoreceptor differentiation, while *Tcf7l1/2* instead promotes RPC maintenance and Müller glia specification. This analysis also highlighted some of the limitations of our model. While we predicted that *Tbx3* overexpression would promote amacrine and Müller glia specification at the expense of rod photoreceptors, we unexpectedly observed that it also promoted specification of bipolar cells. This may reflect the indirect actions of *Tbx3* target genes, as is observed for NFI factors in retinal progenitors (Fig. 5), or could potentially reflect activation of *Tbx2* target genes, since *Tbx2* is predicted to strongly promote expression of bipolar-specific genes (Table ST12). Likewise, while overexpression of *Insm1* and *Insm2* promote rod differentiation, loss of function of *Insm1/2* delays but ultimately does not disrupt rod specification (Fig. 7C), which in turn may reflect the extensive functional redundancy seen among individual TFs in these densely interconnected cell type-specific GRNs. This functional redundancy may complicate the prediction of the developmental phenotype of gain or loss of function of any individual TF, as may interactions with cofactors or post-translational modifications to TFs. The collection of additional functional data of this sort will help refine the GRN models used here, and improve their predictive accuracy.

The data generated in this study serves as a broadly useful resource for the community for further functional characterization of GRNs controlling retinal neurogenesis and cell fate specification, and may help facilitate and improve strategies for reprogramming of endogenous Müller glia and/or directed differentiation of ES/iPS cells to replace neurons lost due to blinding diseases (Javed and Cayouette, 2017; Lahne et al., 2020; Miltner and La Torre, 2019). Sequential expression of TFs that promote formation of early or late-stage neurogenic RPCs, followed by TFs that drive specification of rods, cones or retinal ganglion cells could provide a robust approach to generate these neurons for therapeutic purposes.

### Data availability

All mouse and human scRNA-seq and scATAC-seq data can be accessed at GEO accession number GSE181251. Interactive displays of all scRNA-seq and scATAC-seq data can be accessed through the St Jude Cloud Visualization Community.

## Acknowledgements

The authors would like to thank Richard Gronostajski for providing *Nfia/b/x* conditional mice, Linda Orzolek and Tyler Creamer at the Hopkins Transcriptomics and Deep Sequencing Core for use of the 10x Genomics Chromium controller and library sequencing; and J. Nathans, H. Jaspers, M. Cayouette, A. Javed, L. Goff, J. Kebschull, W. Yap, and members of the Blackshaw lab for comments on the manuscript. This work was supported by the NIH National Eye Institute grants R01EY020560 and U01EY027267 to SB, EY001765 in support of flow sorting, R01EY029548 and P30EY001765 to J.Q., R00EY027844 to B.S.C; P30 EY002687 in support of confocal imaging. S.B. is supported by a Stein Innovation Award from Research to Prevent Blindness. B.S.C is supported by an unrestricted grant to the Washington University Department of Ophthalmology and Visual Sciences and a Career Development Award from Research to Prevent Blindness.

## Supplemental figure legends

**Supplemental Figure 1: Quality control of scATAC-seq data.**

(A) The number of fragments per cell. Bars (cells) are colored by sample and ordered along the x-axis according to fragment number (high to low). The numbers of cells for each time point that passed QC are indicated on the top.

(B) Fragment size distribution (left) and transcriptional start site enrichment profiles (right) of single-cell ATAC-seq. Lines are colored by sample.

(C) Comparison between aggregated scATAC-seq of primary RPCs and bulk ATAC-seq of Chx10- GFP+ retinal progenitors at E11 and P2 (Stein-O’Brien et al., 2019).

(D) Chromatin accessibility plot for the *Hes5* gene locus, showing the similarity between scATAC-seq data and bulk ATAC-seq data. The samples and data types are indicated on the left.

(E) Heatmap showing the Pearson correlations between gene expression and gene accessibility for each retinal cell type. Cell type identities are indicated on the top (scRNA-seq) and right (scATAC-seq) of the heatmap.

(F) The percentages of HMM regions in the cell-type-specific accessible regions at P0 (left) and P14 (right). Bars are colored by 11 different HMM states (Aldiri et al., 2017).

(G) Examples of HMM tracks and cell type-specific aggregate scATAC-seq signal for five marker gene locus in P14: *Aqp4*, *Tfap2b*, *Opn1sw*, *Rho,* and *Capb5*.

**Supplemental Figure 2: Examples of cell-type-specific regulatory elements and motifs.**

(A) Aggregated accessibility profiles of representative cell-type-specific regulatory elements. Each track shows the aggregated scATAC-seq profile from each cell type. The nearest and positively correlated gene of each region is labeled at the top of the plot. The coordinates of these regions are shown at the bottom.

(B) UMAP projection of the scATAC-seq profile shows the activity of the representative cell-type-specific motifs. Cells are colored by chromVAR z-score. The motif ID is indicated at the top of each plot.

132.

132.

**Supplemental Figure 3: UMAP projection of human retinal scATAC-seq and scRNA-seq data.**

(A) UMAP projection and clustering results of human retinal scATAC-seq from gestational day 53-

(B) UMAP projection and clustering results of human retinal scRNA-seq from gestational day 53-

(C) UMAP projection and clustering results of human retinal and retinal organoid scRNA-seq from culture day 24-postnatal day 8 (Lu et al., 2020).

**Supplemental Figure 4: Analytic flowchart for IReNA v2**.

(A) Workflow of Integrated Regulatory Network Analysis (IReNA v2) integrating scRNA-seq and scATACseq data to reconstruct the gene regulatory network (see Methods for detailed description). ArchR, MACS2, TOBIAS and motifmatchr software packages were used in IReNA v2.

(B) Schematic diagram of the integrating method used in IReNA v2 to identify positive and negative transcriptional regulators controlling progenitor state transitions and cell fate specification (see Methods).

**Supplemental Figure 5: *Nfia/b/x* promote late-stage RPCs temporal identity.**

(A) UMAPs and clustering results of scRNA-seq data and scRNA-seq data. Shading indicates cell

(B) Heatmaps of DEGs in primary RPCs/MG . Each row represents a DEG, and each column represents a cell (left). The DEGs are ordered by their log2 fold change (treatment/control) as shown in the middle panel. The distributions of cell-type-specific genes among DEGs were shown on the right panel.

(C) Heatmaps of CARs in primary RPCs/MG. Each row represents a DAR, and each column represents a different condition (left). The DARs are ordered by their log2 fold change (treatment/control) as shown in the middle panel. The distributions of cell-type-specific peaks among DARs were shown on the right panel. The top5 enriched motifs are listed on the right.

(D) Heatmaps showing *Nfia/b/x* ChIP-seq signal at P2. Around 13,680 peaks were identified.

(E) Comparison of *Nfia/b/x* binding regions with gene annotation and accessible regions in P2.

(F) Bar plot showing that *Nfia/b/x* binding regions are strongly enriched in stage 3 RPC-specific accessible regions.

(G) Venn diagrams showing the overlaps between predicted *Nfia/b/x* regulated genes and cell-type-specific DEGs. The p-value on the top of each Venn diagram indicates the significance of their overlap as determined by the hypergeometric test.

**Supplemental Figure 6: Gene regulatory networks controlling specification of retinal neurons.**

(A,E) Models of retinal cell states during intermediate (A) and late stages of retinal neurogenesis (E). UMAPs of scRNA-seq and scATAC-seq data from E18-P2 retina (A) and P5-P8 retina (E). Shading indicates cell type.

(B,F) UMAPs showing differentiation trajectories inferred from scRNA-seq and scATAC-seq at intermediate (B: E18-P2) and late stages (F: P5-P8) of retinal neurogenesis. Shading indicates pseudotime status.

(C,G) Heatmaps showing the expression of cell type-specific DEGs (left) and the accessibility of their corresponding CARs (right) along these differentiation trajectories. The top bars are colored by pseudotime state for each trajectory.The left bar indicates cell types and the classes of CARs .

(D,H) Networks showing feedback relationships among TF pairs selectively expressed in primary and neurogenic RPCs, as well as postmitotic neurons (left). Each node represents an individual TF. Each edge represents a positive or negative feedback regulatory relationship between TF pairs.

**Supplemental Figure 7: Identification of transcription factors controlling neurogenesis in postnatal retina.**

(A) Candidate TFs predicted to control rod specification inferred from E18-P2 GRNs, rank ordered by the number of cell type-specific TFs and non-TF genes predicted to be directly regulated by each TF. Gene names outlined in red indicate regulatory relationships previously validated using genetic analysis.

(B) Candidate TFs predicted to maintain RPC cell status inferred from the E18-P2 GRNs.

(C) UMAP plots showing *Insm1/2*, *Tbx3*, and *Tcf7l1/2* expression at E18-P2 in mouse retina (left). UMAPs showing *INSM1/2*, *TBX3*, and *TCF7L1/2* expression at gestational week 14-20 in the human retina (right).

(D) UMAP plots of scRNA-seq data from following gain and loss of function analysis of candidate TFs.

## Supplemental Tables

**Table ST1:** Number of cells of each type identified at each developmental age using scATAC-Seq in mouse retina.

**Table ST2:** Cell-type specific cis-regulatory elements and transcription factor motifs identified in developing mouse retina.

**Table ST3:** Number of cells of each type identified at each developmental age using scATAC-Seq in human retina.

**Table ST4:** Cell-type specific cis-regulatory elements and transcription factor motifs identified in developing human retina.

**Table ST5:** Evolutionarily conserved cell-type specific cis-regulatory elements and transcription factor motifs identified in developing retina.

**Table ST6:** Differentially expressed genes (DEGs) and correlated differentially-accessible chromatin regions (CARs) in stage 1-3 RPCs and Müller glia.

**Table ST7:** Regulatory relationships among transcription factors that comprise cell type-specific gene regulatory networks in stage 1-3 RPCs and Müller glia.

**Table ST8:** Differentially expressed genes (DEGs) and differentially-accessible chromatin regions (DARs) seen in RPCs and Müller glia following overexpression or knockdown of *Nfia/b/x*.

**Table ST9:** Location of Nfia/b/x ChIP-Seq peaks in P2 retina.

**Table ST10:** Differentially expressed genes (DEGs) and correlated differentially-accessible chromatin regions (CARs) in neurogenic RPCs and subtypes of developing retinal neurons.

**Table ST11:** Regulatory relationships among transcription factors that comprise cell type-specific gene regulatory networks in neurogenic RPCs and subtypes of developing retinal neurons.

**Table ST12:** List of transcription factors that are predicted to activate and/or repress expression of genes specific to neurogenic RPCs and subtypes of developing retinal neurons.

**Table ST13:** Differentially expressed genes (DEGs) observed following overexpression or knockdown of *Insm1/2*, listed by cell type.

**Table ST14:** Differentially expressed genes (DEGs) observed following overexpression or knockdown of *Tcf7l1/2* or *Tbx3*, listed by cell type.

## Methods

### Animals and retinal cell dissociation

CD1 mice were purchased from Charles River Laboratories. All experimental procedures were pre-approved by the Institutional Animal Care and Use Committee (IACUC) of the Johns Hopkins University School of Medicine. Mouse embryos or pups at different timepoints of retinal development (E11, E12, E14, E16, E18, P0, P2, P5, P8, P11 and P14) were used for this study. *Chx10-Cre-EGFP;Nfia^fl/fl^;Nfib^fl/fl^;Nfix^fl/fl^* mice were generated as described previously (Clark et al., 2019). Mice were euthanized, and eyes were removed and incubated in ice-cold PBS. Retinas were dissected, and cells were dissociated using Papain Dissociation System as described previously (Hoang et al., 2020). Each sample contains a minimum of 4 retinas from 4 animals, regardless of sex. Dissociated cells were resuspended in ice-cold PBS containing 0.04% bovine serum albumin (BSA). Cell count and viability were assessed by Trypan blue staining.

### *Ex vivo* retinal electroporation and fluorescence-activated cell sorting (FACS)

Retinas from CD1 mouse embryos at day 14 (E14) and postnatal day 0 (P0) were used for *ex vivo* electroporation as described previously (de Melo and Blackshaw, 2011). For overexpression studies, pCAGIG was used as a control, while pCAGIG-based plasmids encoding full-length ORFs were used for overexpression (see Table). For analysis of *NFIA/B/X* and TCF7L1/2 function, equal molar amounts of each plasmid were combined prior to electroporation.

For somatic Crispr-mediated gene knockout, the *CBh* promoter of Cas9-P2A-GFP plasmid (Addgene #48138) was replaced by pCAG promoter (pCAGIG, Addgene #11159) to allow for more robust Cas9-P2A-GFP expression in retinal explants. Dual gRNAs targeting two different exon regions were cloned into a single Cas9 plasmid using PrecisionX™ Multiplex gRNA Cloning Kit with U6 and H1 promoters. gRNAs were designed using CHOPCHOP tool (https://chopchop.cbu.uib.no/). For combined *Insm1/2* knockout, equal molar amounts of each gRNA-Cas9 plasmid were mixed prior to electroporation. Retinal cells were dissociated from explants for fluorescence-activated cell sorting (FACS) as described previously (Hoang et al., 2020). GFP+ cells were collected in ice-cold PBS with 10% heat-inactivated fetal bovine serum (FBS). To determine Crispr-mediated knockout efficiency, genomic DNA was extracted from GFP+ cells from *Insm1/2* knockout and empty Cas9 control, and subjected for PCR and digestion using GeneArt™ Genomic Cleavage Detection Kit.

### Immunohistochemistry

Explants used for immunohistochemical analyses were cultured to P0 or P11 equivalent (6 or 11 days *in vitro*), fixed in 4% paraformaldehyde in PBS, and processed through sucrose gradients before mounting in OCT compound, cryosectioning (15 µm sections), and immunohistochemical analyses. Stained slides of retinal explant sections were imaged using a Zeiss LSM 800 confocal microscope. For each immunostaining condition, 2-3 single-plane confocal images per retinal explant were counted, with counts aggregated across individual explants. Individual data points shown in Figures 5D and 7C represent cell counts obtained from individual explants.

### Single Cell RNA-seq library construction and sequencing

ScRNA-seq analysis was performed on dissociated retinal cells using 10x Genomics. Briefly, dissociated retinal cells (∼10,000 cells per sample) were loaded into a 10x Genomics Chromium Single Cell system using Chromium Single Cell 3’ Reagents Kits v3.1 (10X Genomics, Pleasanton, CA). scRNA libraries were generated by following the manufacturer’s instructions. Libraries were pooled and sequenced on Illumina NextSeq 500 or NovaSeq 6000. Sequencing reads were processed through the Cell Ranger 3.1 pipeline (10x Genomics) using default parameters.

### Single Cell ATAC-seq library construction and sequencing

ScATAC-seq was performed using the 10x Genomic single cell ATAC reagent v1.1 kit following the manufacturer’s instruction. Briefly, dissociated cells were centrifuged at 300xg for 5 min at 4°C. Cell pellet was resuspended in 100 μl of Lysis buffer, mixed 10x by pipetting and incubated on ice for 3 min. Wash buffer (1 ml) was added to the lysed cells, and cell nuclei were centrifuged at 500xg for 5 min at 4^0^C. Nuclei pellet was re-suspended in 250 μl of 1x Nuclei buffer. Cell nuclei were then counted using Trypan blue. Re- suspended cell nuclei (10-15k) were used for transposition and loaded into the 10x Genomics Chromium Single Cell system. Libraries were amplified with 10 PCR cycles and were sequenced on Illumina NextSeq or NovaSeq with ∼200 million reads per library. Sequencing data was processed through the Cell Ranger ATAC 1.1.0 pipeline (10x Genomics) using default parameters.

### ChIP-seq

Freshly dissected P2 retinas were homogenized and cross-linked for 10 minutes using 1% formaldehyde (ThermoFisher Scientific Cat# 28906) on a tube rotator at room temperature. Glycine was added to a final concentration of 0.125 M to quench the cross-linking reaction and washed three times with ice-cold PBS with cOmplete protease inhibitors (Millipore Sigma Cat# 11836170001). The cells were then prepared for sonication using the truChIP chromatin shearing kit (Covaris Cat# 520154). Briefly, cells were lysed at 4C on a tube rotator for 10 minutes using the 1X lysis buffer B. Intact nuclei were then collected by centrifugation at 1700xg for 5 minutes and washed with 1X wash buffer C before being resuspended in 1ml of 1X shearing buffer D3. Nuclei were then transferred to 1mL milliTUBE with AFA fiber (Covaris Cat# 520130) and sonicated using the E220 focused-ultrasonicator (Covaris Cat# 500239). Chromatin immunoprecipitation was then performed on the sheared DNA using the iDeal ChIP-seq kit for transcription factors (Diagenode Cat# C01010170). One percent of chromatin was kept aside to be used as an input control. Antibodies against targets used for chromatin immunoprecipitation are NFI (NFIA, NFIB, NFIX), and IgG. Briefly, equal volume of sheared chromatin was incubated overnight with 3 μg of antibody in iC1b buffer with protease inhibitors and BSA and washed DiaMag Protein A-coated magnetic beads (Diagenode Cat# C03010020-220) on a tube rotator at 4°C overnight. The magnetic beads were then washed sequentially with wash buffers iW1, iW2, iW3 and iW4. DNA was then de-crosslinked and eluted for 4 hours at 65C before being purified using IPure beads (Diagenode Cat# C03010014). The purified DNA was then subjected to sequencing library preparation or qPCR analysis. Libraries were prepared from 5ng of DNA using the Ovation Ultralow System V2 (Tecan Genomics Cat# 0344NB-32) and sequenced on the Illumina NextSeq500.

### Single-cell ATAC-seq analysis

#### Preprocessing

The Cell Ranger (Zheng et al., 2017) ATAC pipeline was used to process the raw sequencing data for mapping, de-duplication and identification of Tn5 cut sites. We first convert BCL files to fastq format with the function ‘cellranger-atac mkfastq’. Then, we mapped the fastq files to the mm10 genome (refdata-cellranger-atac-GRCh38-1.2.0) with the function ‘cellranger-atac count’. This function outputs the aligned, barcoded, and Tn5 insertion corrected fragment files, which were used for all downstream analysis.

#### Filtering cells by TSS enrichment, unique fragments, nucleosome banding and doublet score

The ArchR package(Granja et al., 2020) was used to process the fragment files. We calculated the TSS enrichment, unique fragments and nucleosome banding for each cell with the function ‘createArrowFiles’’. Then we kept the high-quality cells with the following criteria: 1) The number of unique nuclear fragments > 1000. 2) TSS enrichment score > 10. 3) nucleosome banding score < 4. We next identified potential doublets with the function ‘addDoubletScores’, and removed doublets using ‘filterDoublets’’ with the following parameters: cutEnrich = 2, cutScore = -Inf, and filterRatio =2. Finally, we filtered the fragment files according to the cells we retained. These cleaned fragment files were used for all downstream analysis.

#### Generating union peaks

Union peaks were generated for all the samples as described by Satpathy, et al. (Satpathy et al., 2019). We first constructed 2.5kb tiled windows across the mm10 genome and computed a cell-by-window sparse matrix by counting Tn5 insertion (from cleaned fragment files) overlaps for each cell. Next, we binarized the cell-by-window matrix and created a Seurat object for each sample with the Signac (Stuart et al., 2020) package. Then we performed dimension reduction and clustering analysis using the functions ‘RunTFIDF’, ‘RunSVD’, ‘FindNeighbors’ and ‘FindClusters’ with 2-50 dimensions and 0.3 resolution. Next, we call peaks for each identified cluster in each sample using MACS2 (Zhang et al., 2008) software with the following parameters: ‘-shift -75 --extsize 150 --nomodel --callsummits --nolambda --keep-dup all -q 0.05’.

We further extended the peak summits on both sides to a final width of 500 bp, and filtered these fixed-width peaks if they overlapped with mm10 v2 blacklist regions. (https://github.com/Boyle-Lab/Blacklist/blob/master/lists/mm10blacklist.v2.bed.gz). Finally, we kept the top 100,000 fixed-width peaks for each cluster in each sample according to their -log10(q-value) and then merged them to the final union peak sets using the ‘reduce’ function from the GenomicRanges (Lawrence et al., 2013) package.

### LSI clustering, visualization, and identification of cell types

For each sample, the cell-by-peak matrix was generated by the union peak sets, and was binarized and inputted to the Signac pipeline. Then we performed dimension reduction, clustering and UMAPs analysis under the standard Signac workflow.

To annotate the cell types for each cluster, we used existing mouse and human scRNA-seq datasets (Clark et al., 2019; Thomas et al., 2021) to interpret our scATAC-seq cell types using the CCA (canonical correlation analysis) integration method in the Seurat package. Firstly, we downloaded the mouse scRNA-seq data (“https://github.com/gofflab/developing_mouse_retina_scRNASeq”) and converted them to Seurat objects. Secondly, for each scATAC-seq sample, we calculate the ‘gene activity’ profile for each cell with the function ‘CreateGeneActivityMatrix’. Finally, for each age-matched sample pair from scATAC-seq and scRNA-seq datasets, we identify anchors between them with the function ‘transfer.anchors.’, and we used the ’TransferData’ to obtain the cell type prediction results for each cell. We further filtered out cells with a prediction score < 0.5 and annotated each cluster according to their predicted cell types.

### Integration of E11-P14 single-cell ATAC-seq datasets

To Integrate and visualize all the cells from the scATAC-seq data (E11 to P14), we used the following 3 steps:

1. Filtering cell types. To better focus on the epigenetic difference during the retinal development, we removed the cells which are not annotated as retinal cells in each time point before **integration**. We kept the following cell types: RPCs, Neurogenic, RGC, AC/HC, Cone, Rod, BC and MG. The total cell-by-peak matrix is filtered according to the retinal cells and used in the downstream analysis.
2. Selecting variable peaks. To remove the potential batch effect, we selected the variable peaks separately in each sample. Because scATAC-seq data is very sparse, we aim to aggregate similar cells to create a more dense cell-by-peaks matrix to facilitate variable peaks calling. First, based on the UMAP embedding, we used the kNN approach to find the 100 nearest cells for each individual cell. Then we aggregate raw counts for each cell by its corresponding 100 nearest cells to create a new cell-by-peaks aggregate matrix. We then identified the variable peaks based on the new matrix using Seurat pipelines: ‘NormalizeData’ and ‘’FindVariableFeatures’ (selection.method = “mvp”). Finally, we combined all the variable peaks from each sample into a master variable feature set, which was used in the downstream dimension reduction and clustering procedure.
3. LSI clustering and visualization. Firstly, we binarized the filtered cell-by-peak matrix from Step1 and performed the TF-IDF normalization. Then we used the master variable feature set from Step 2 to perform the dimension reduction with ‘RunSVD.’ Next, we used the 2nd-20th dimensions to identify clusters with a resolution of 1, and calculated the UMAP coordinates for visualization. Finally, we plotted the 3D UMAP of all retinal cells with the plotly graphing library in Python.

### ChromVAR and footprint analysis

We used the chromVAR (Schep et al., 2017) R package to infer global TF activity in each cell.

Firstly, we fed the total raw cell-by-peak matrix into chromVAR and to correct for GC bias with the mm10 reference genome. Next, we generated a TF z-score matrix with the mouse TF Motif database (TransFac2018) using the function ‘computeDeviations’. The z-score for each cell was used to generate the heatmap and visualization using previously calculated UMAP coordinates.

To analyze and plot TF footprints in different retinal cell types, we used the same methods described in Corces et al. (Corces et al., 2018). First, we predicted the TF binding sites with the TF PWM matrix and the identified accessibility region using the function ‘matchmotifs’ in motifmatchr R package. Second, we generated 3 tables: 1) Table1: An aggregated observed 6-bp hexamer table in ± 250bp region relative to all the motif centers. 2) Table2: An aggregated expected 6-bp hexamer table from the mm10 genome. 3) Table3: An observed Tn5 insertion signal table around the ± 250bp relative to the motif centers. Next, we obtain the O/E 6-bp hexamer table by dividing the two hexamer tables (Table5 = Table1/Table2). Then we normalized the signal using the O/E 6-bp hexamer table (Table4/ Table5) to get the final Tn5 bias-corrected signal.

### Single-cell RNA-seq analysis

#### Preprocessing

We processed raw scRNA-seq data with the Cell Ranger software for formatting reads, demultiplexing samples, genomic alignment, and generating the cell-by-gene count matrix. The ‘cellranger mkfastq’ function was used to convert BCL files to fastq files. The ‘cellranger count’ function was used to process fastq files for each sample with the mm10 mouse reference index provided by 10x Genomics. The cell-by-gene count matrix is the final output from the Cell Ranger pipeline. We used the cell-by-gene count matrix for all downstream analysis.

We applied the Seurat (Stuart et al., 2019) package to create Seurat objects for each sample with the cell-by-gene count matrix and the function ‘CreateSeuratObject’ (min.cells = 3, min.features = 200). After visual checking the violin plot of the total counts for each cell, we filtered out cells with nCount_RNA < 800 or nCount_RNA > 8000. We further filtered out the cells with a mitochondrial fraction > 8%. Next, we used Scrublet (Wolock et al., 2019) to identify multiplet artifacts, and removed potential doublet cells from each sample using default parameters. Non-neuroretinal cell types, such as microglia and astrocytes, were also removed.

#### Trajectory inference and pseudotime analysis

We applied Slingshot (Street et al., 2018) software to infer trajectories based on the UMAP coordinates. We only kept the cells involved in the developmental process we plan to investigate. Then we ran Slingshot with the UMAP coordinates matrix and set ‘RPCS1’ cluster (Figure4) and ‘RPC’ cluster (Figure6,FigureS6) as the root cells to calculate the trajectory with the function “getLineages” and “getCurves”. Then, we applied the “slingPseudotime ’’ function to calculate pseudotime state for each cell. Finally, we calculated the bin-by-gene matrix or bin-by-peak matrix by averaging expression levels or accessibility levels of all the cells in each bin.

#### Identification of cell-type specific peaks and motif activities

We calculated the cell-type specific peaks across all the retina cells for mouse and human scATACseq respectively. The function ‘getMarkerFeatures’ in ArchR package were used to identify the marker peaks for each cell type with the following parameters: normBy=’nFrags’, bias=c(“TSSEnrichment”,”log10(nFrags)”) and testMethod = “wilcoxon”. We then further filtered the results to get the final specific peak sets with the function “getMarkers”: cutOff = “FDR <= 0.01 & Log2FC >= 1.5”.

The cell-type-specific motifs were identified based on the chromVAR results. Firstly, we identify the significant enriched motif for each cell type. The chromVAR deviation matrix was converted to a Seurat object. Then the cell type information were added to the Seurat object and the enriched motif for each cell type was measured by the function “FindAllMarkers” with following parameters: only.pos = TRUE, test.use = ’LR’, and p value < 0.01. Secondly, we further filtered the significant motifs according to their average Z-score and their ranks among the retina cell types. We only kept the motifs for each cell type if they 1) average chromVAR Z-score > 1, and 2) their average chromVAR Z-score are the highest or the second highest among all the cell types.

#### Identification of conserved peaks between mouse and human

We compared cell-type specific peaks between mouse (mm10) and human (hg38) by the rtracklayer package in R. we converted mouse peak region from mm10 assembly to hg38 assembly with the function ‘liftOver’. Then we identified the overlapped peak pairs between mouse converted peaks and human peaks with the function ‘findOverlaps’. We also calculated the overlapped ratio for each pair as: Overlapped ratio = width(Mouse converted peak) / width(Overlapped region). Finally, we identified the pairs of peaks as conserved peak pairs if their overlapped ratio > 0.5.

#### Gene/peaks set enrichment analysis

We performed Gene/peak set enrichment analysis using the fgsea package in R using default parameters (Korotkevich et al., 2021). The significant DEGs and DARs were ranked based on their log2 fold change (treatment / control). The retina cell-type-specific gene sets and peak sets we used in the GSEA / PSEA were generated from all the retinal cells as mentioned before.

#### Constructing gene regulatory networks by integrating scRNA-seq and scATAC-seq data

To infer cell-type-specific GRNs from scRNA-seq and scATAC-seq data, we modified IReNA pipeline to IReNA v2, which contains the following main modules:

1. Selecting candidate genes The DEGs were used as candidate genes for GRNs construction. For each developmental process we aim to investigate, we identified the enriched genes for each cell type using the function ‘FindMarkers’ in Seurat. In constructing the GRNs of progenitors transition, the following parameters of ‘FindMarkers’ were used: min.pct = 0.05, logfc.threshold = 0.20, only.pos = TRUE, p-adjust < 0.01. In constructing GRNs regulating neurogenesis, the following parameters of ‘FindMarkers’ were used: min.pct = 0.1, logfc.threshold = 0.25, only.pos = TRUE and p-adjust < 0.01.
2. Identifying significant peak-to-gene links We used the ArchR package to identify the significant peak-to-gene links. First, we integrated the age-matched scRNA-seq and scATAC-seq datasets for each time point using unconstrained Integration method with the function ‘addGeneIntegrationMatrix’. Then, using the function ‘addPeak2GeneLinks’, we calculated the correlation between accessibility peak intensity and gene expression. Finally, we identified the significant peak-to-gene links with the following cutoff: abs(correlation) > 0.2 and fdr < 1e-6.
3. Identifying the potential cis-regulatory elements for each candidate gene We identified potential cis-regulatory elements for each candidate gene based on their location and the peak-to-gene links from Step2. We first classified all peaks into three categories according to their genomic location related to their potential target genes: 1) Promoter. 2) Gene body. 3) Intergenic. For the peaks in the promoter region,we treated all of them as correlated differentially-accessible chromatin regions (CARs) of their target genes. For the peaks in the gene body region, we defined them as CARs if they met the following criteria: 1) the distance between the peak and the TSS of its target gene is < 100kb. 2) the links between the peak and its target gene is significant. For the peaks in the intergenic region, we first find their target genes and construct the peak-gene pairs if the target genes’ TSS are located within the upstream 100kb or downstream 100 kb of the intergenic peaks. Then we keep the peak-gene pairs if their peak-to-gene links are significant in step2. These peaks were identified as CARs of their gene pairs.
4. Predicting cell-type specific TFs binding in cis-regulatory elements With the cis-regulatory elements identified in Step 3, we next predicted the TF binding in these elements for each cell type with the PWMs extracted from TRANSFAC database. Firstly, we searching the motifs in all the cis-regulatory elements with the function ‘matchMotifs (p.cutoff = 5e-05)’ from the motifmatchr package. Then we filtered these motif regions according to their footprint score and their corresponding TF’s expression for each cell type. To calculate the footprint score for each motif region in each cell type, we re-grouped the insertion fragments based on their origin of cell type and converted these cell-type-specific fragments into bam files using a custom script. Then we fed the bam files to TOBIAS software and obtained the bias-corrected Tn5 signal (log2(obs/exp)) with the default parameters except: ATACorrect --read_shift 0 0. Next, we calculated footprint scores including NC, NL and NR for each motif’s binding region. NC indicated the average bias-corrected Tn5 signal in the center of the motif. NL and NR indicated the average bias-corrected Tn5 signal in the left and right flanking regions of the motif, respectively. The flanking region is triple the size of the center region. We kept the motifs with the following criteria: NC < -0.1 and NL > 0.1 and NR > 0.1. We further removed the motifs binding region for each cell type if the expression level of their corresponding TFs are not enriched in that cell type (from Step1).
5. Calculating gene-gene correlation We calculated the expression correlations between all the expressed genes at the single-cell level. First, we extracted the cell-by-matrix from Seurat objects and filtered out the non-expressed genes in the matrix (rowSums < 10). Then we applied the MAGIC software to impute missing values and recover the gene interactions using the cell-by-gene matrix. The output matrix from MAGIC was used to calculate gene-gene correlation using the function ‘cor’ in R. To identify the significant gene-gene correlations, we ranked all the gene-gene correlations (∼1X10e8). The top 2.5% correlations were treated as significant positive correlations (p < 0.025) and the bottom 2.5% correlations were treated as significant negative correlations (p < 0.025).
6. Constructing gene regulatory networks By integrating data from Step1-Step5, We constructed cell-type specific GRNs with the following procedure: We first obtained the peak-target links from Step 3, and cell-type specific TF-peak links from Step 4. We then merged these 2 types of links to the cell-type specific TF-peak-target relationships. Next, we classified these TF-peak-target relationships into activation or repression relationships based on the sign of the expression correlation between TF and target from Step 5. The significant positive/negative correlated TF-targets were selected as the active/repressive regulations respectively. Finally, we removed all the duplicated TF-target regulatory relationships for each cell type and merged them to the final GRNs which were used for the downstream analysis.
7. Identifying and visualizing feedback TF pairs With the GRNs constructed in the previous steps, we searched for TF pairs connected by either positive or negative feedback regulatory relationships. The TF pairs that activated each other were identified as ‘double positive’ pairs and the TF pairs repressed each other were identified as ‘double negative’ pairs. We visualized these feedback TFs pairs using Cytoscape software.

### Constructing inter-cell type regulatory networks2

We used a previously described approach (Hoang et al., 2020) to calculate the significance of regulatory relationships between cell types with the constructed GRNs. For each cell type, we first selected their highly enriched genes with the cutoff: q <0.01 and logFC > 0.5. Then we calculated the number of active and repressive regulations between these highly enriched genes for each cell type from the GRNs and calculated p-value with the hypergeometric test. We set p-value < 0.01 as a cutoff to determine the significant regulations.

### ChIP-seq data analysis

After removing adaptors with Trimmomatic (Bolger et al., 2014), we mapped the cleaned fastq files to mm10 genome using bowtie2 (Langmead and Salzberg, 2012). We next filtered low quality reads with SAMtools (Li et al., 2009)(MAPQ < 10), and removed PCR duplicates using Picard tools (http://broadinstitute.github.io/picard/) . For NFI ChIP-seq data, we used IgG and Input samples as control, and used MACS2 (Zhang et al., 2008) to call peaks with the default parameters except: -q 0.01. Finally, we identified 13,680 NFI binding peaks and used them in the downstream analysis.

### Integration analysis of single-cell RNA-seq or single-cell ATAC-seq datasets between control and treatment samples

We applied the Harmony (Korsunsky et al., 2019) package to integrate the scATAC-seq data from different genotypes at the same age (control vs NFIA/B/X overexpression, or control vs *Nfia/b/x* knockout). Briefly, we first merged the cell-by-peak matrix from the same age, then inputted the cell-by-peak matrix into the Signac analysis pipeline. We normalized and obtained a low-dimensional representation of the cell-by-peak matrix using the functions ‘FindTopFeatures’, ‘RunTFIDF’ and ‘RunSVD’. Next, we integrated all the cells from different genotypes (Ctrl vs NFI Overexpress or Ctrl vs NFI TKO) using the ‘RunHarmony’ function with the options: dim.use = 2:50, group.by.vars = ‘genotypes’, reduction = ’lsi’ and project.dim = FALSE. Finally, we used the harmony dimensions to identify clusters, and calculated UMAP coordinates for visualization.

### Inferring Nfia/b/x targets in progenitors

We predicted NFI target genes by integrating the information obtained from scRNA-seq, scATAC-seq and ChIP-seq analysis using the following steps:

1. Identify DEGs resulting from *NFIA/B/X* overexpression and *Nfia/b/x* conditional knockout. We performed the differential gene expression analy sis between control RPC/MG cells and *Nfia/b/x* TKO or *NFIA/B/X* overexpressing RPC/MG cells using the function ‘FindMarkers’ with the options: min.pct = 0.1, logfc.threshold = 0.25. Then we selected the DEGs with adjusted p-value < 0.01.
2. Identify differential peaks following *NFIA/B/X* overexpression or *Nfia/b/x* conditional knockout. To explore which ATAC regions are changed following *NFIA/B/X* overexpression or *Nfia/b/x* conditional knockout, we applied the MAnorm (Shao et al., 2012) algorithm to perform the differential peak analysis between control and *NFIA/B/X* overexpression or *Nfia/b/x* conditional knockout. First, we selected cells in the ‘RPC’ and ‘MG’ cluster and then separated these cells according to their genotypes (control, *NFIA/B/X* overexpression or *Nfia/b/x* conditional knockout). Next, we aggregated the cells in the same condition by summing their count signals for each peak, and created a new condition-by-peak count matrix, and fed this into the MAnorm pipeline. We performed the MAnorm test and identified the differential peaks using the cutoff: LOG_P > 5, abs (M_value_rescaled) > 0.5 and A_value_scaled > 4.
3. Predict NFIA/B/X binding sites in RPC With the same method in the GRN construction, we first identify Peaks-Target links according to peak location and “peak-to-gene links” (identified in RPC-MG developmental process), these cis-regulatory elements including TSS peaks, correlated gene body peaks and intergenic peaks. We further filtered these PeaksTarget links with the following criteria: 1) Peaks should overlap with NFI ChIP-seq peaks. 2) Gene body peaks and intergenic peaks should be differentially accessible following *NFIA/B/X* overexpression or *Nfia/b/x* conditional knockout. 3) Target gene should be differential following *NFIA/B/X* overexpression or *Nfia/b/x* conditional knockout.

### GO term analysis

To understand what biological functions are enriched in the gene set we are interested, we applied Gorilla (Eden et al., 2009) algorithm to calculate the enriched Gene Ontology terms for our gene sets with the default parameters (P-value threshold = 0.001, ontology = ‘Process’).

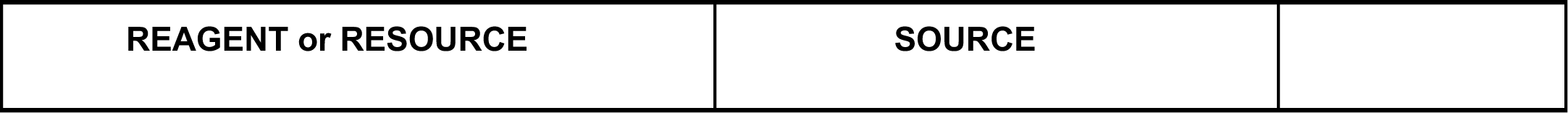
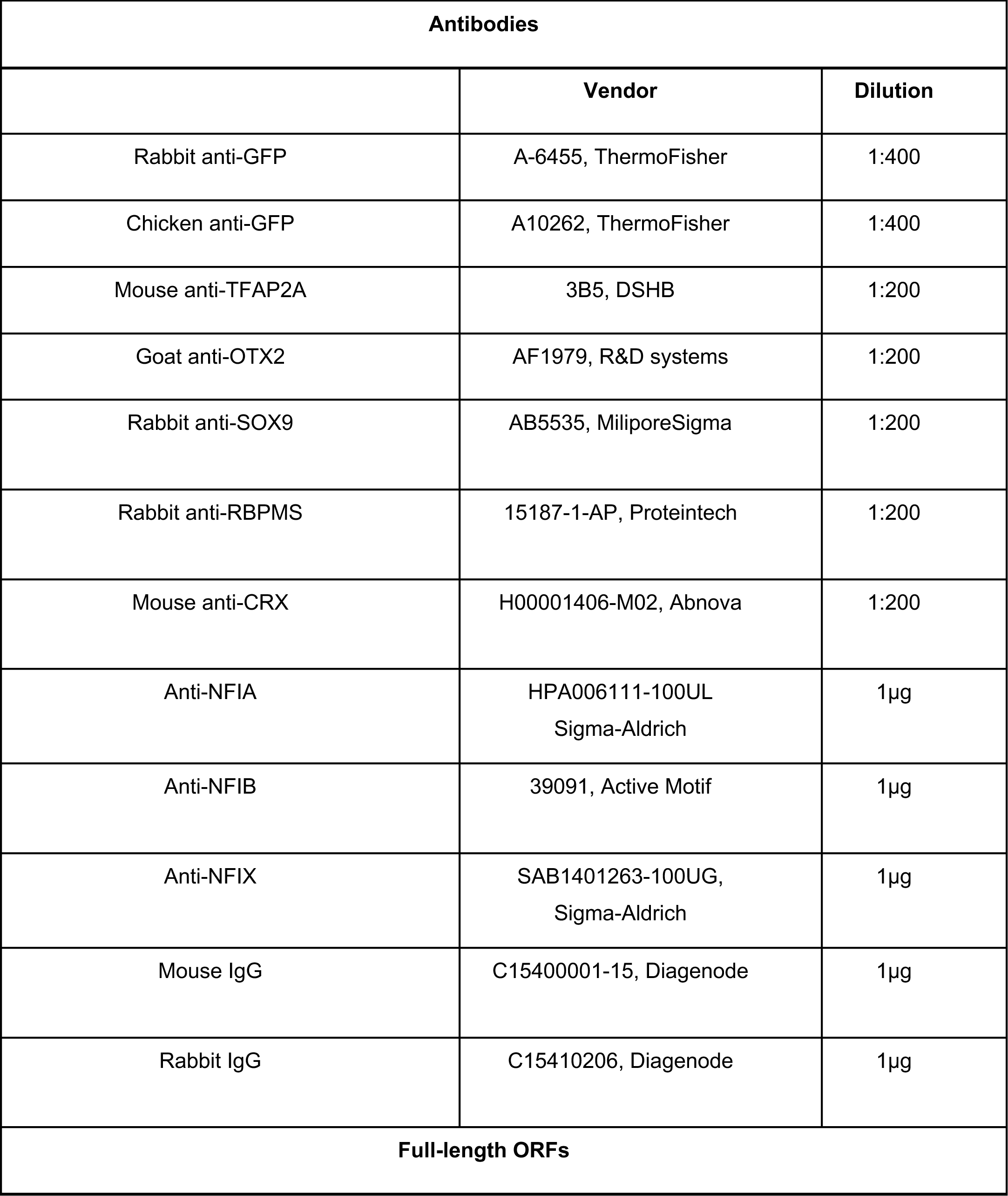
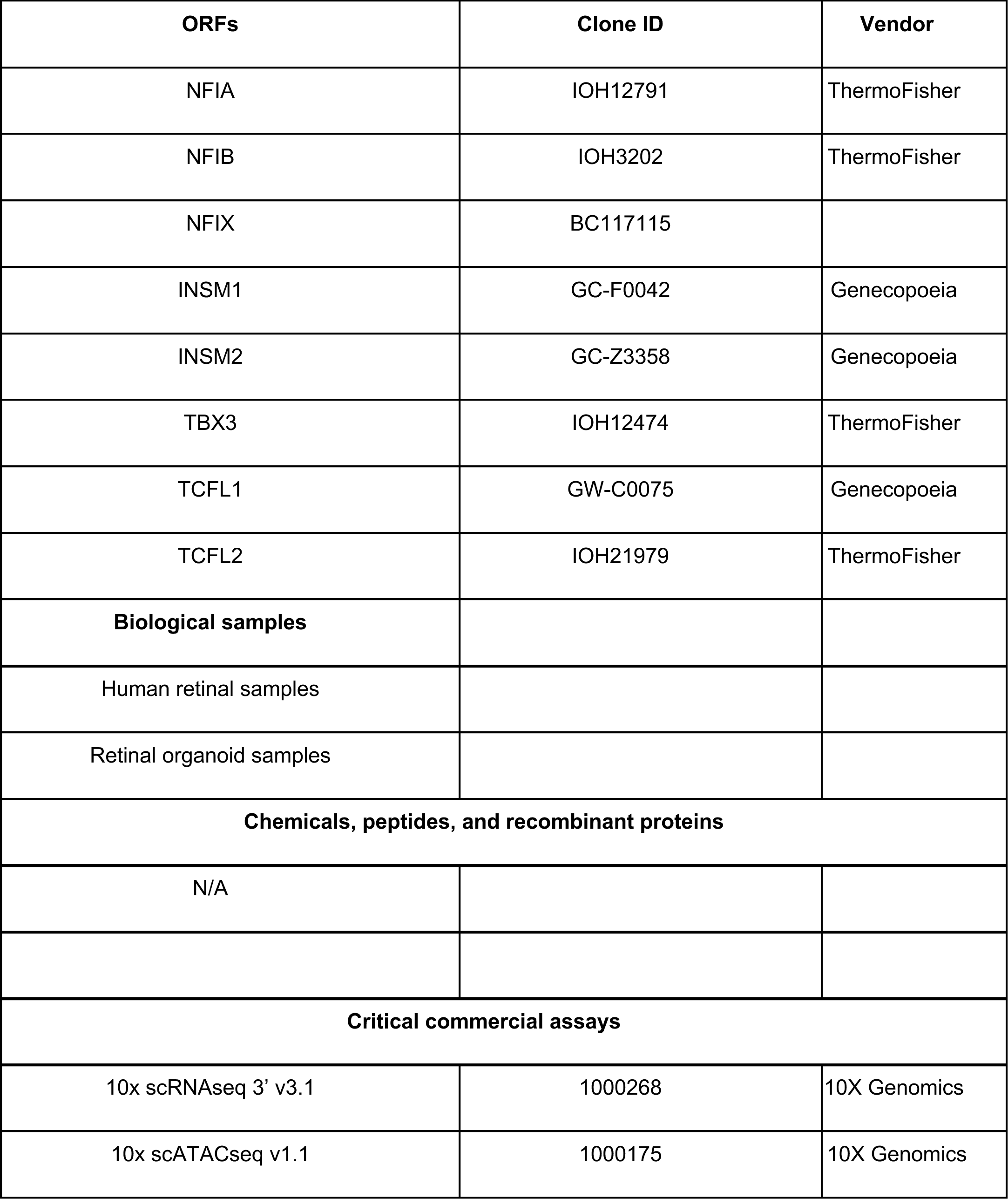
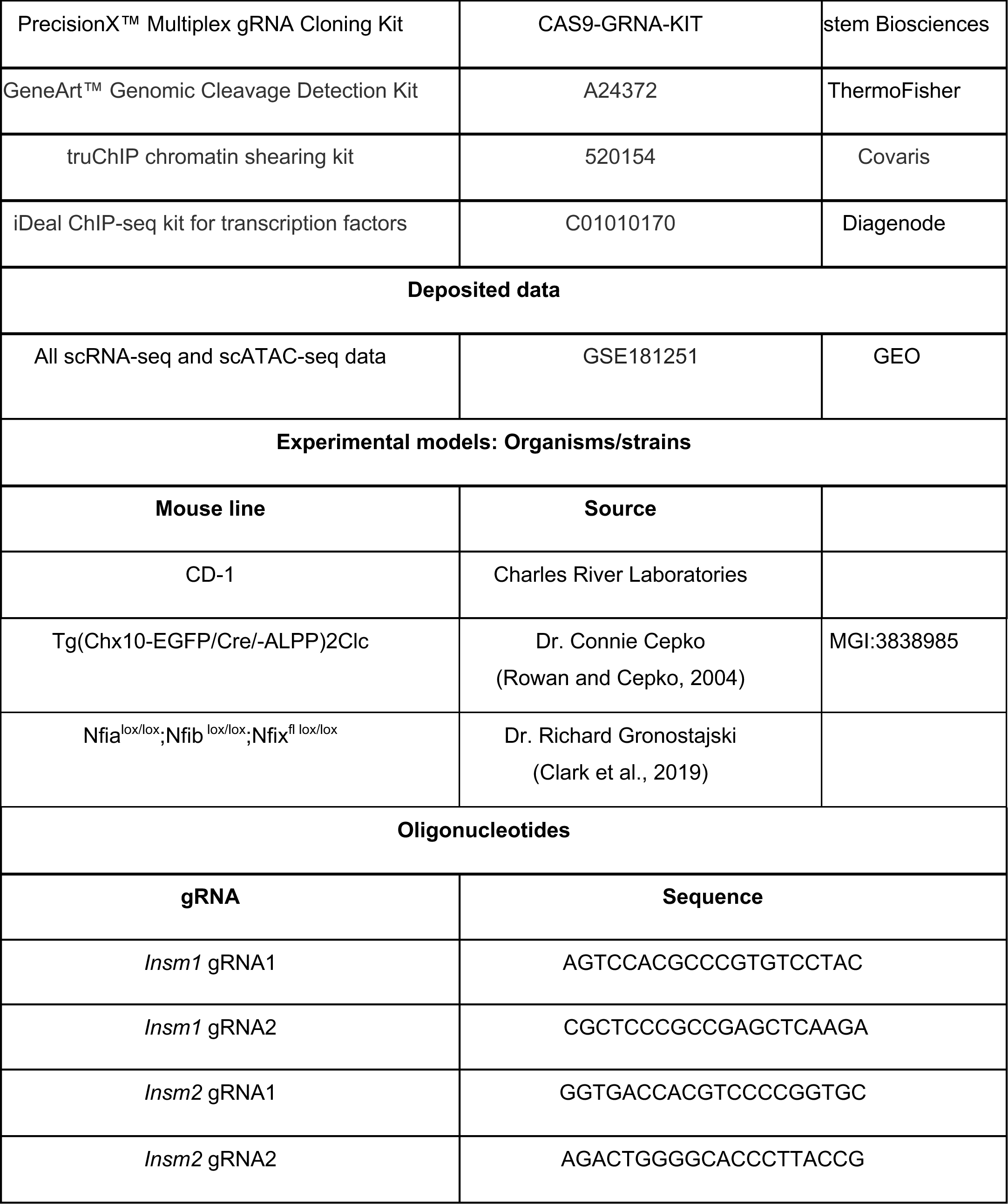
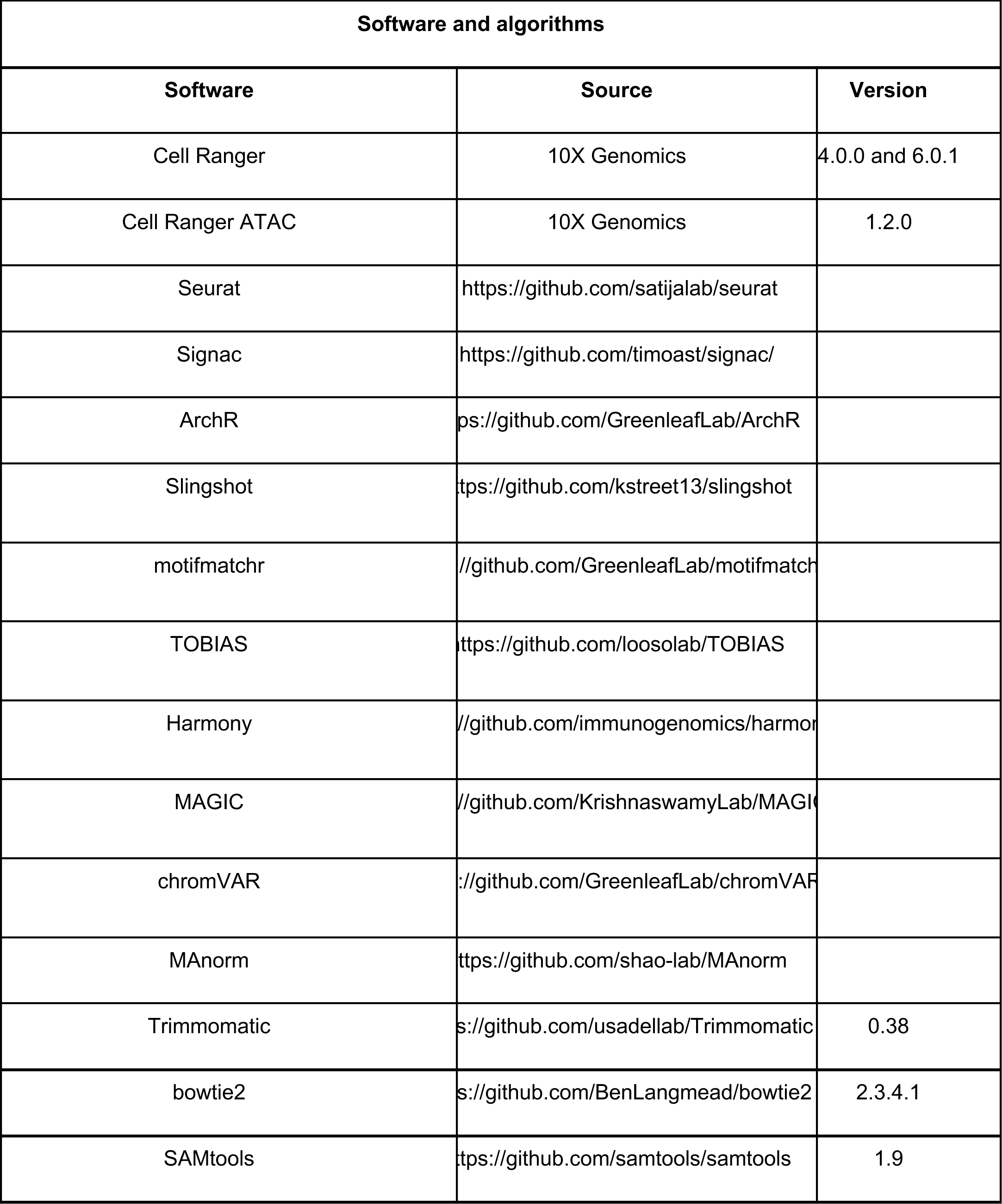
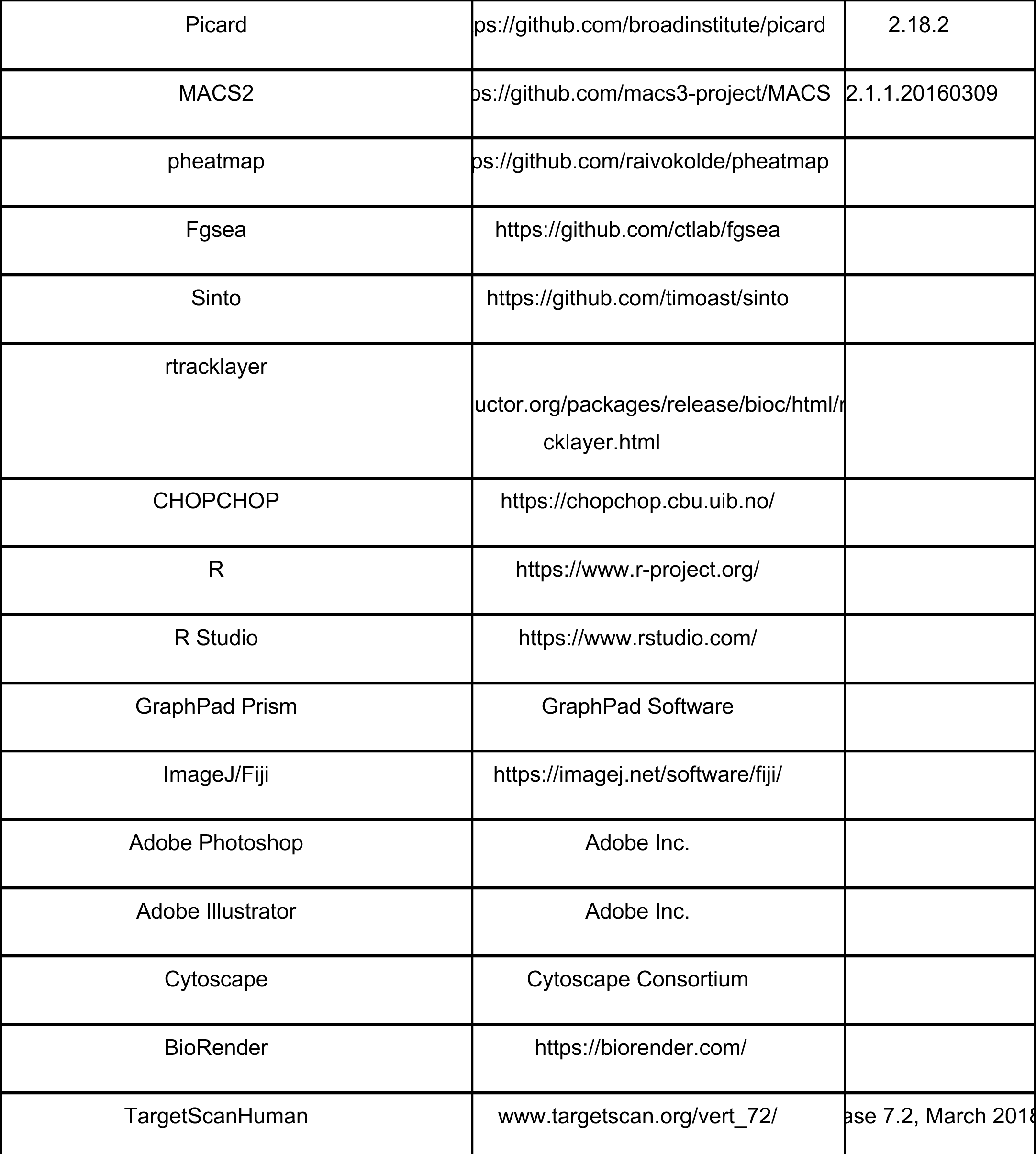
KEY RESOURCES TABLE.

