## Supplementary figures and images for "Gene regulatory networks controlling temporal patterning, neurogenesis, and cell fate specification in the mammalian retina"

### Supplemental Figures S1-S7

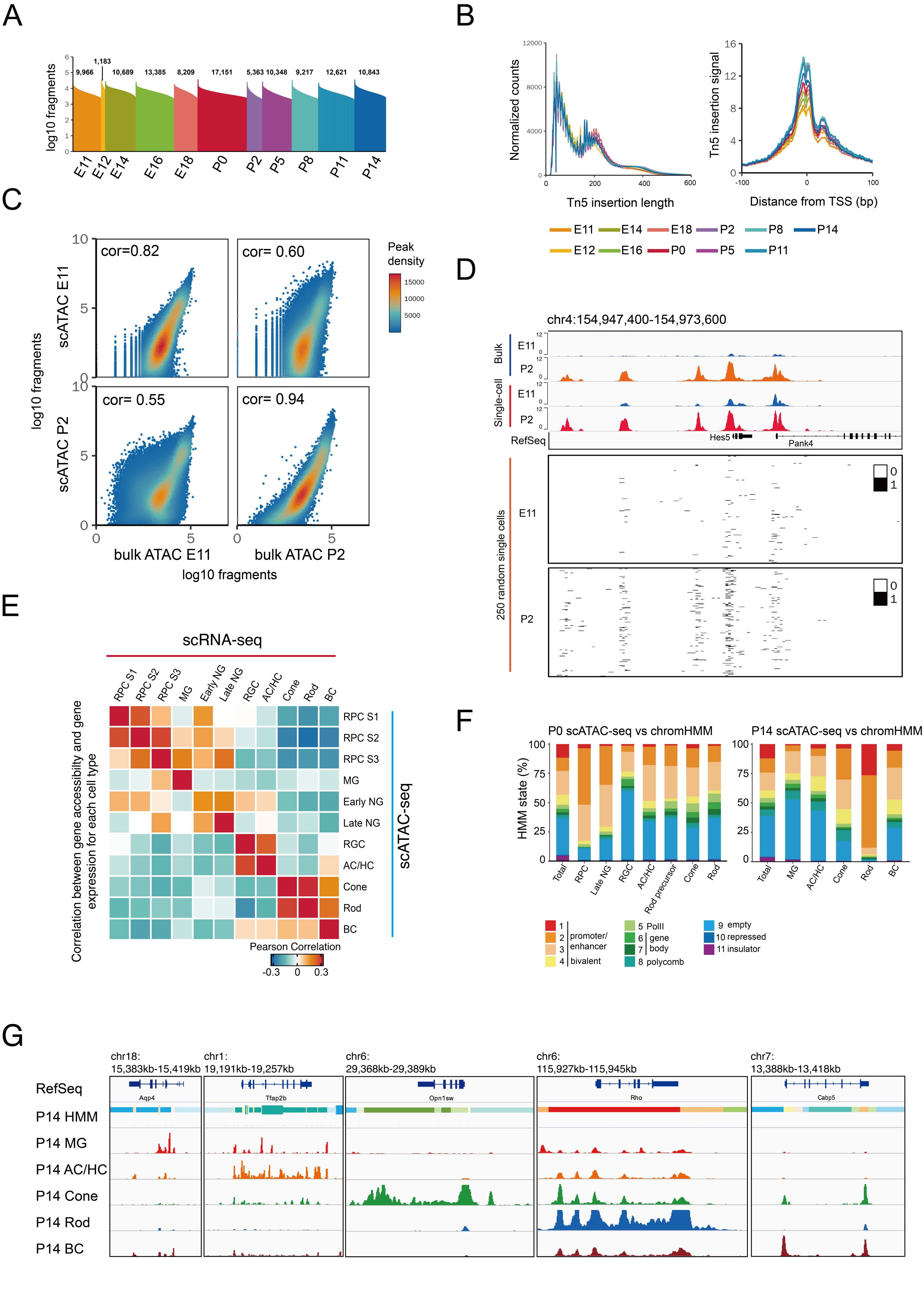


Lyu, et al. Fig.S1

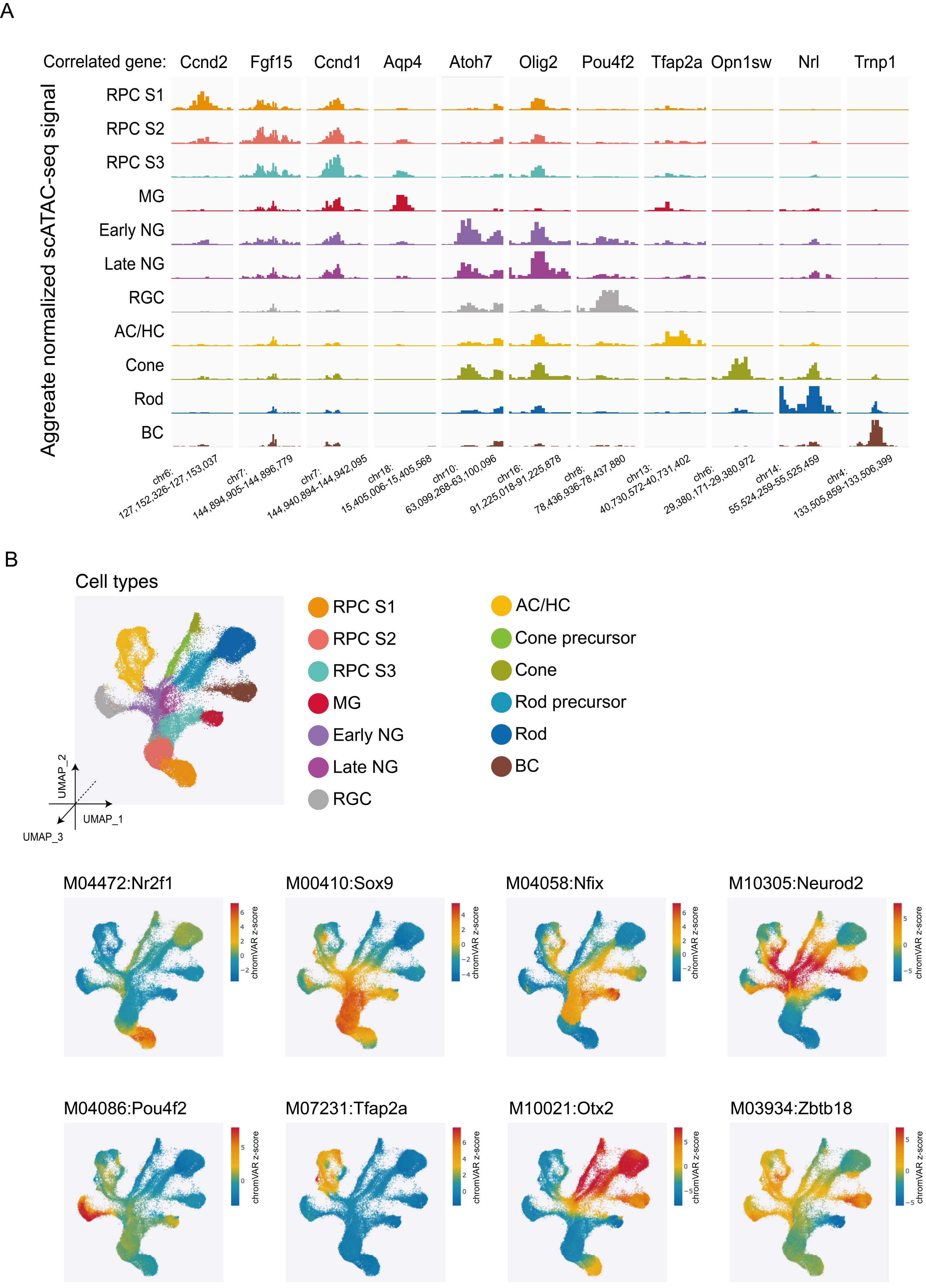


Lyu, et al. Fig.S2

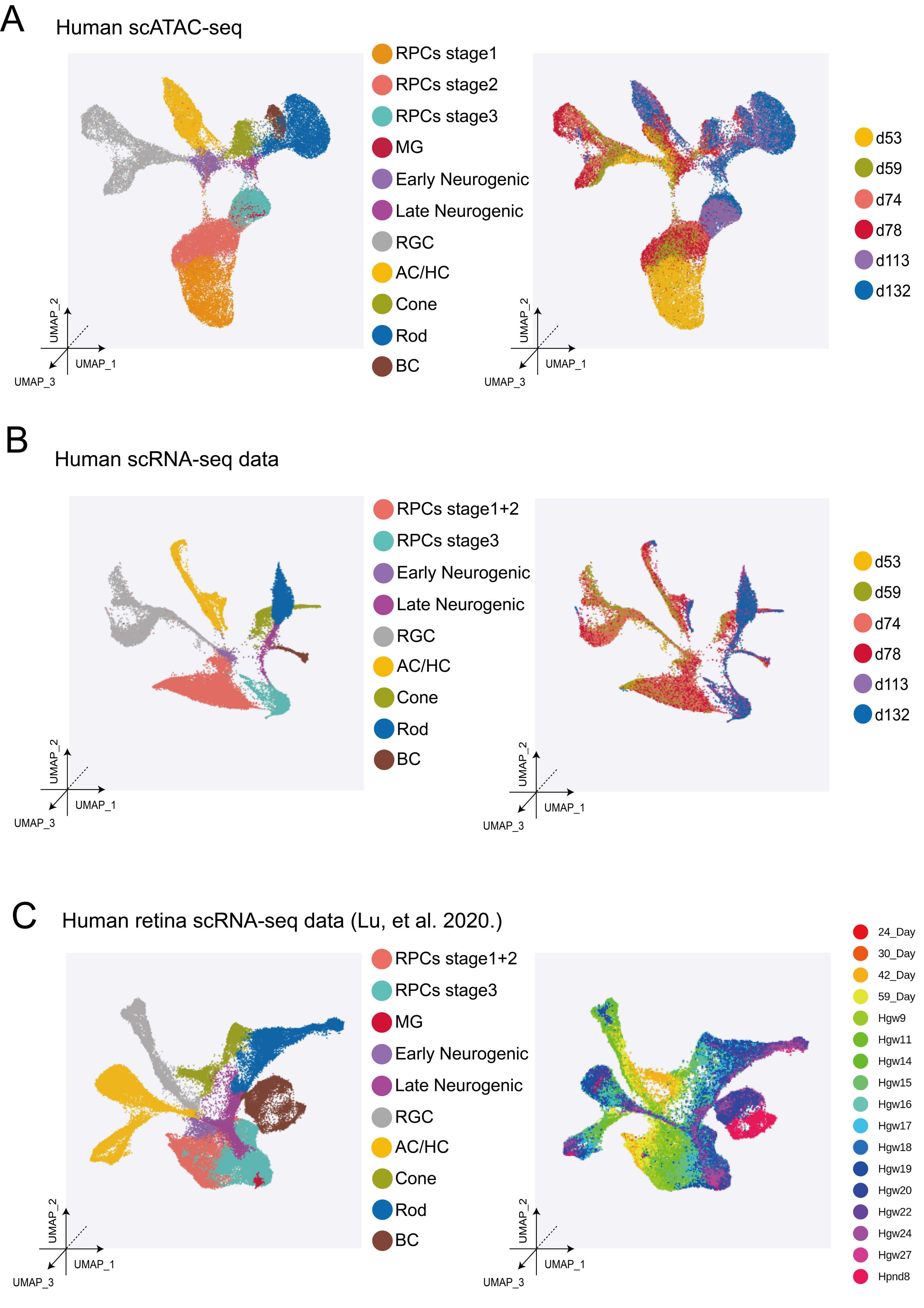


Lyu, et al. Fig.S3

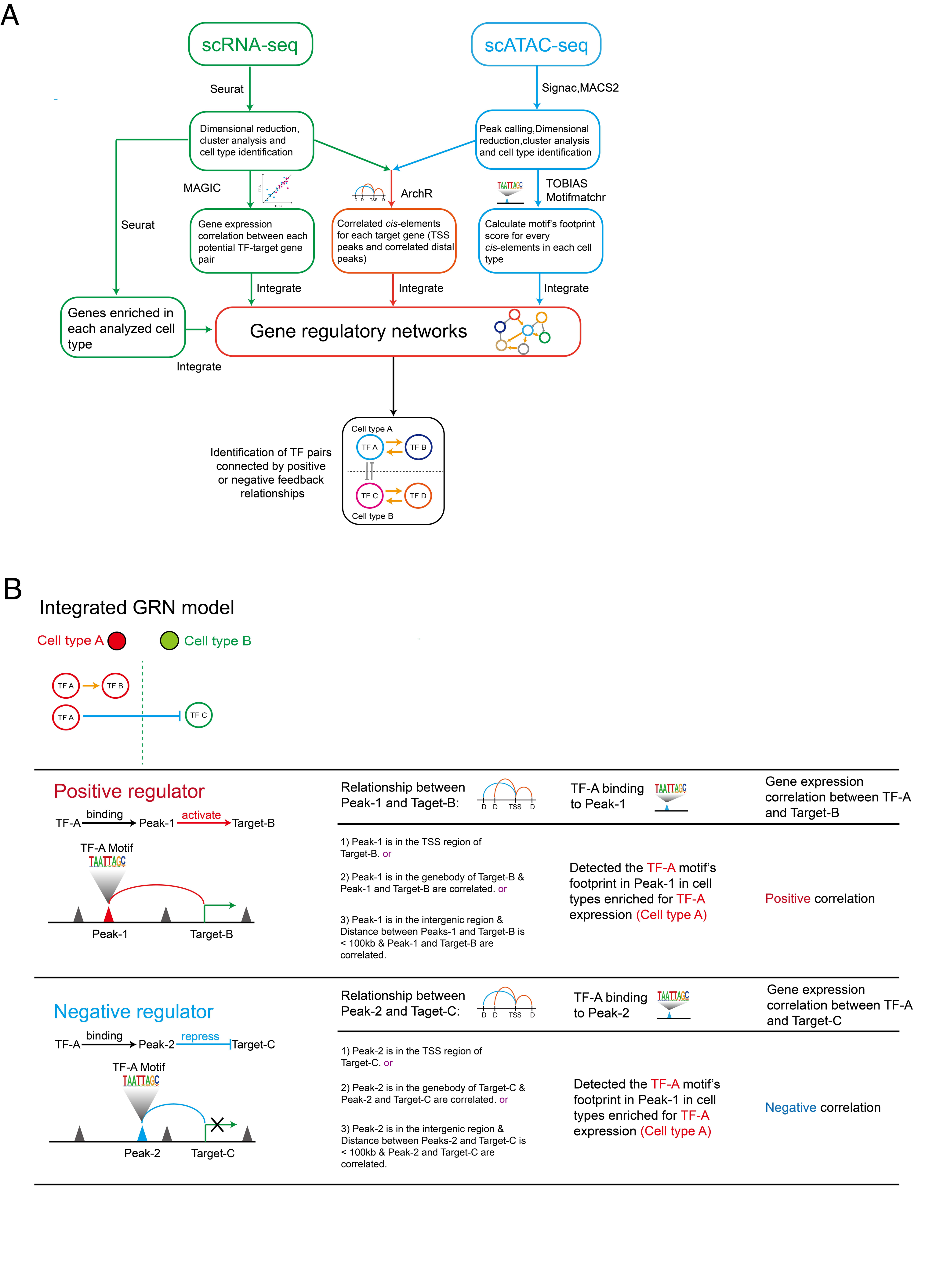


Lyu, et al. Fig. S4

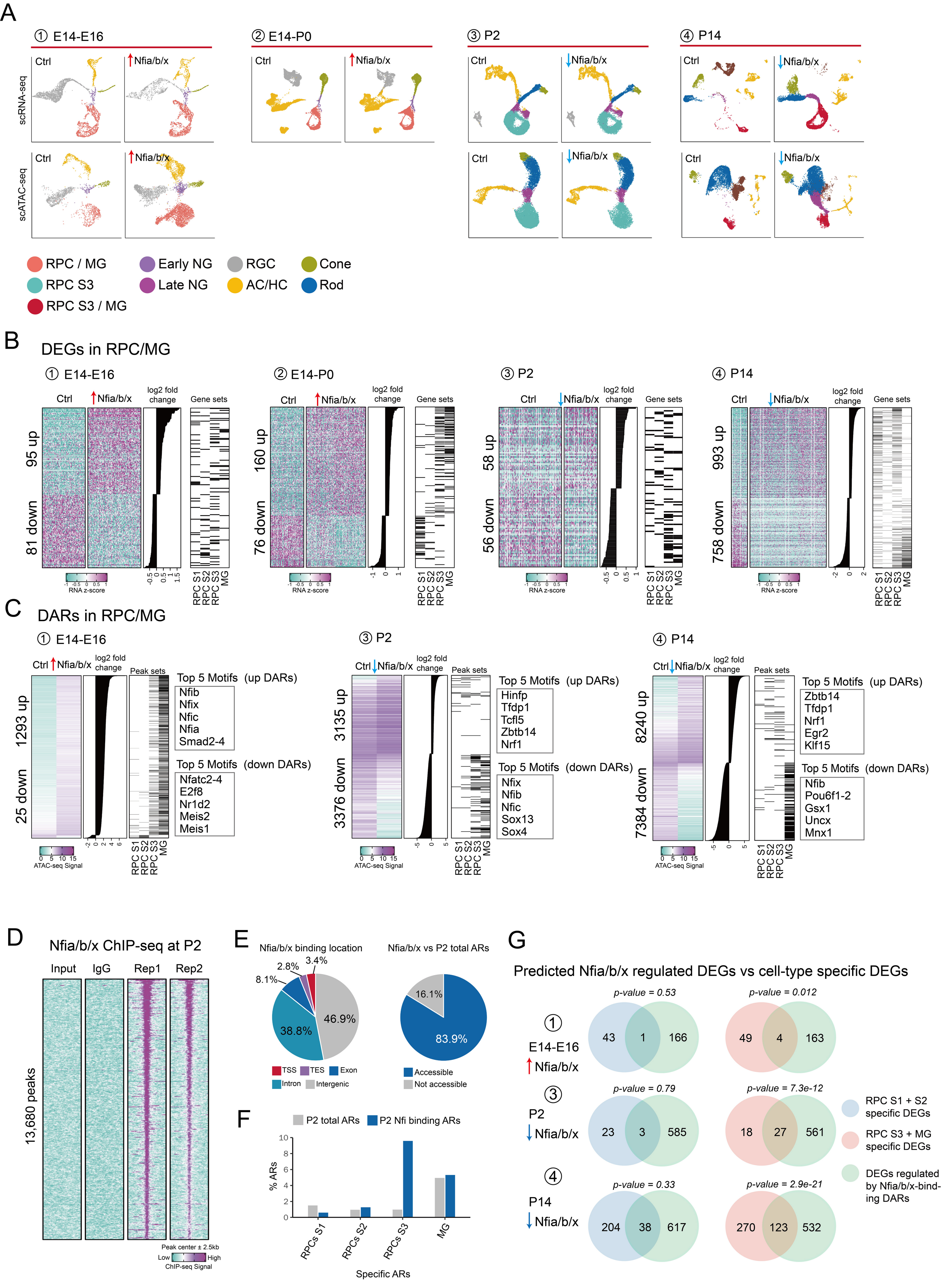


Lyu, et al. Fig. S5

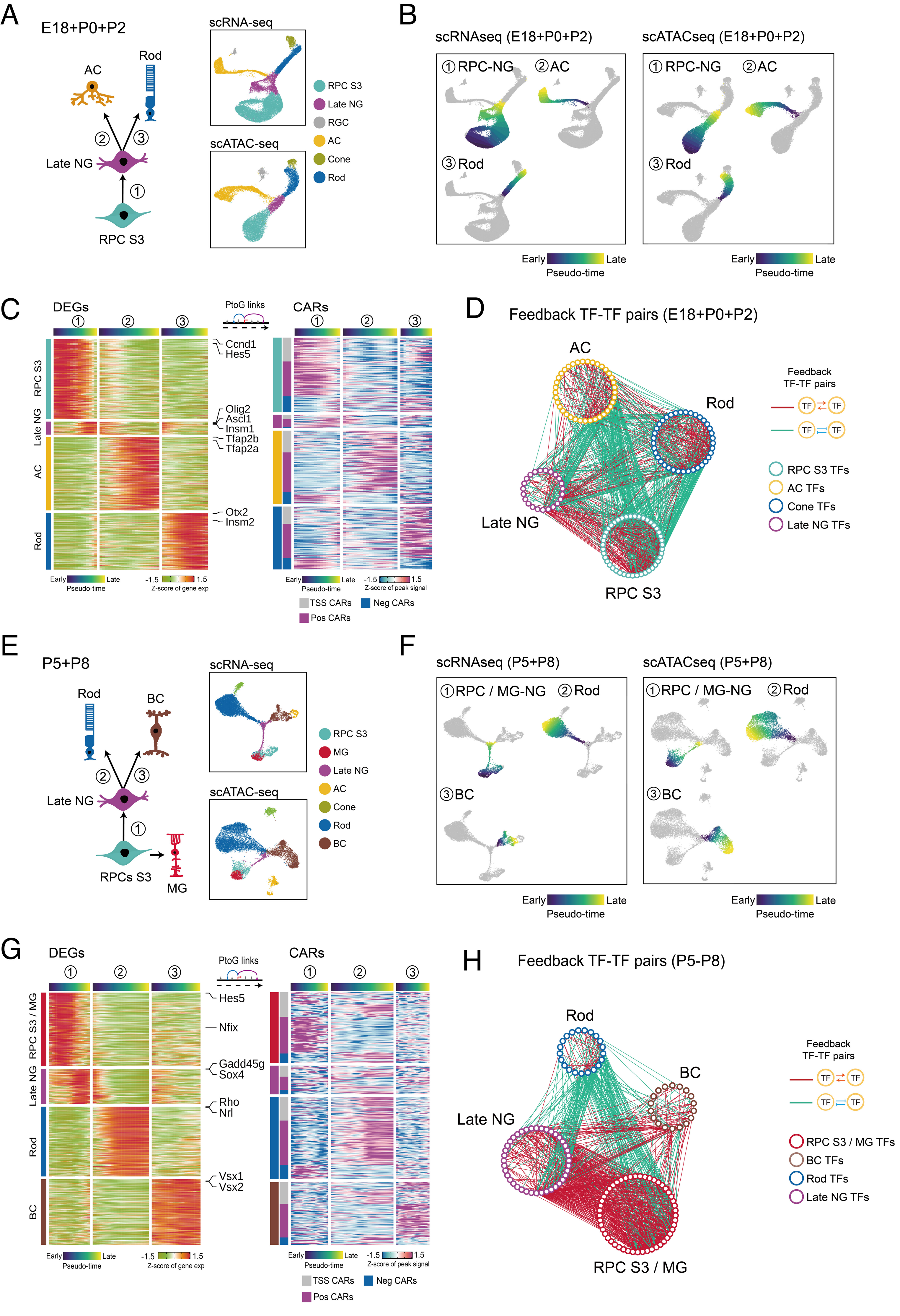


Lyu, et al. Fig. S6

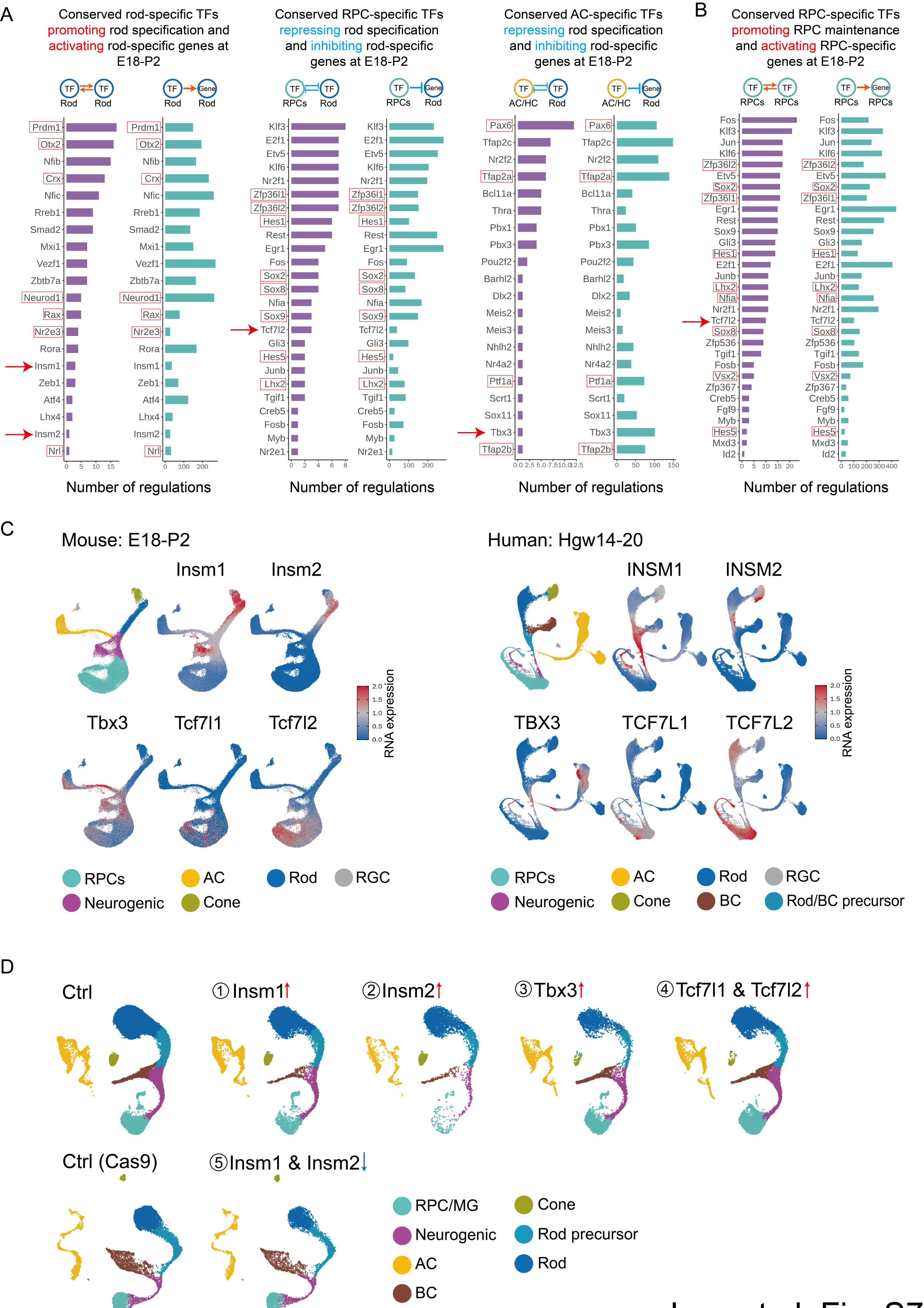


Lyu, et al. Fig. S7
